# *The entire brain, more or less, is at work*: ‘Language regions’ are artefacts of averaging

**DOI:** 10.1101/2023.09.01.555886

**Authors:** Sarah Aliko, Melissa Franch, Viktor Kewenig, Bangjie Wang, Greg Cooper, Annette Glotfelty, Benjamin Hayden, Steven L Small, Jeremy I Skipper

## Abstract

Models of the neurobiology of language suggest that a small number of anatomically fixed brain regions are responsible for language functioning. This derives from centuries of aphasia studies and decades of neuroimaging. The latter rely on thresholded measures of central tendency applied to activity patterns from heterogeneous stimuli. We hypothesised that these methods obscure the whole brain distribution of regions supporting language. Specifically, ‘language regions’ are connectivity hubs, coordinating varying peripheral activity that averages out following thresholding. We tested this with neuroimaging meta-analyses and movie-fMRI. Results show that words localise to ‘language regions’ when averaged but are distributed throughout the brain when examining specific linguistic representations. These ‘language regions’ are partially connectivity hubs that are spatiotemporally dynamic, making connections with ∼40% of the brain periphery outside those regions, and only appearing in the aggregate over time. Hub-periphery connections encode linguistic representations, not the ‘language regions’ alone. Results replicate with audiobook-fMRI. Finally, intracranial neuronal recordings support findings, showing ‘language regions’ are hub-like, with linguistic representations decodable throughout the brain. Together, these four studies suggest that ‘language regions’ are artefacts of averaging heterogeneous language representations. Instead, they are connectivity hubs coordinating whole-brain distributed network representations, suggesting why their damage results in aphasia.

---

> *There is no ‘centre of Speech’ in the brain any more than there is a faculty of Speech in the mind. The entire brain, more or less, is at work in a [person] who uses language*.
>
> — William James^1^

> *Beware of the Blob! It creeps and leaps and glides and slides..*.
>
> — Burt Bacharach and Mack David^2^

## Introduction

Historical views of ‘the organisation of language and the brain’ derive from mid-19th century post-mortem lesion analyses. For the next 100 years, there existed an ongoing debate as to whether language is localised to specific brain regions or holistic, involving the whole brain. The holistic view lost ground in the 1960s when Norman Geschwind helped swing the pendulum back to a localizationist model.^3^ Roughly, this model posits that a left posterior superior temporal lobe (or ‘Wernicke’s area’; but see^4^) and left inferior frontal gyrus (or ‘Broca’s area’) are the anatomical loci for language comprehension and speech production, respectively.^5^ These localisations and ‘localizationism’ have subsequently remained the standard view,^6^ likely for several reasons.^7^ Most prominently, beginning in the 1970s and continuing until now, tens of thousands of *in vivo* positron emission tomography (PET) and functional magnetic resonance imaging (fMRI) studies of language appear to support these anatomical observations.

Advances from neuroimaging have arguably been only marginally incremental over Geschwind’s now ‘classical’ model. Specifically, contemporary models of the systems neurobiology of language recognise that ‘language’ is a complex process that can be decomposed into subprocesses that are localised to a few additional regions. These regions are all in close proximity to those in the classical model or their right hemisphere ‘homologues’ but not typically ascribed to that older model. They include more of the superior temporal lobe, as anterior as the temporal pole, middle temporal lobe, and premotor cortices.^6,8–11^ This neo-localizationist view is communicated in the scientific literature using phrases like ‘language regions’ to describe portions of the brain and ‘the language network’^1^ to describe commonly co-active regions of the brain.^6,7^ Indeed, these conceptualizations seem to be validated by the anatomical regions emerging from neuroimaging meta-analyses, conducted across a wide variety of language stimuli and tasks (see Fig. 2).

**Table 1.**
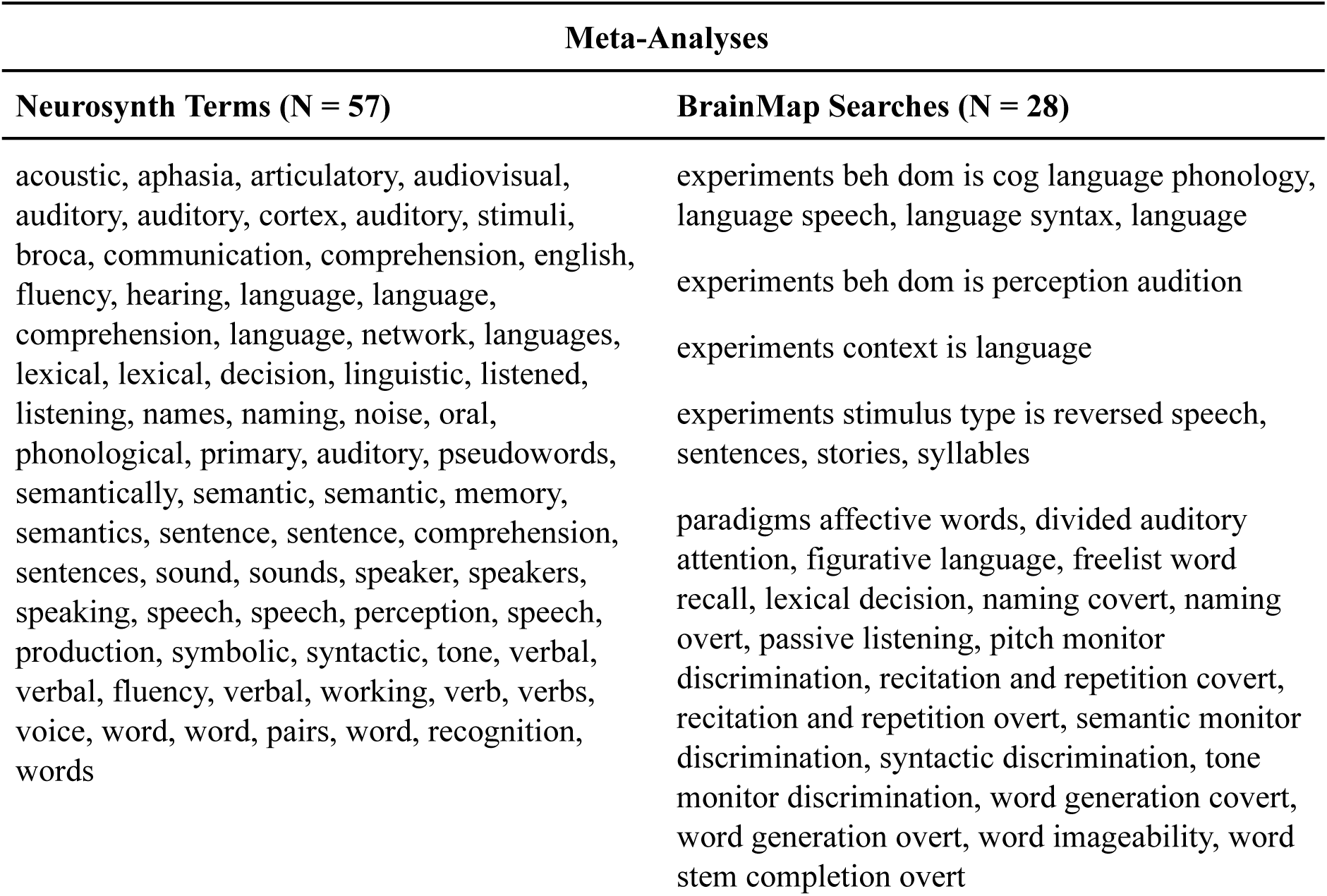
Terms and searches used to conduct the meta-meta-analysis. Each Neurosynth term in the left-hand column corresponds to a single meta-analysis. Each row in the right-hand column lists the Sleuth search criteria used to identify studies for one or more meta-analyses in BrainMap (see Fig. 2 for results).

**Table 2.**
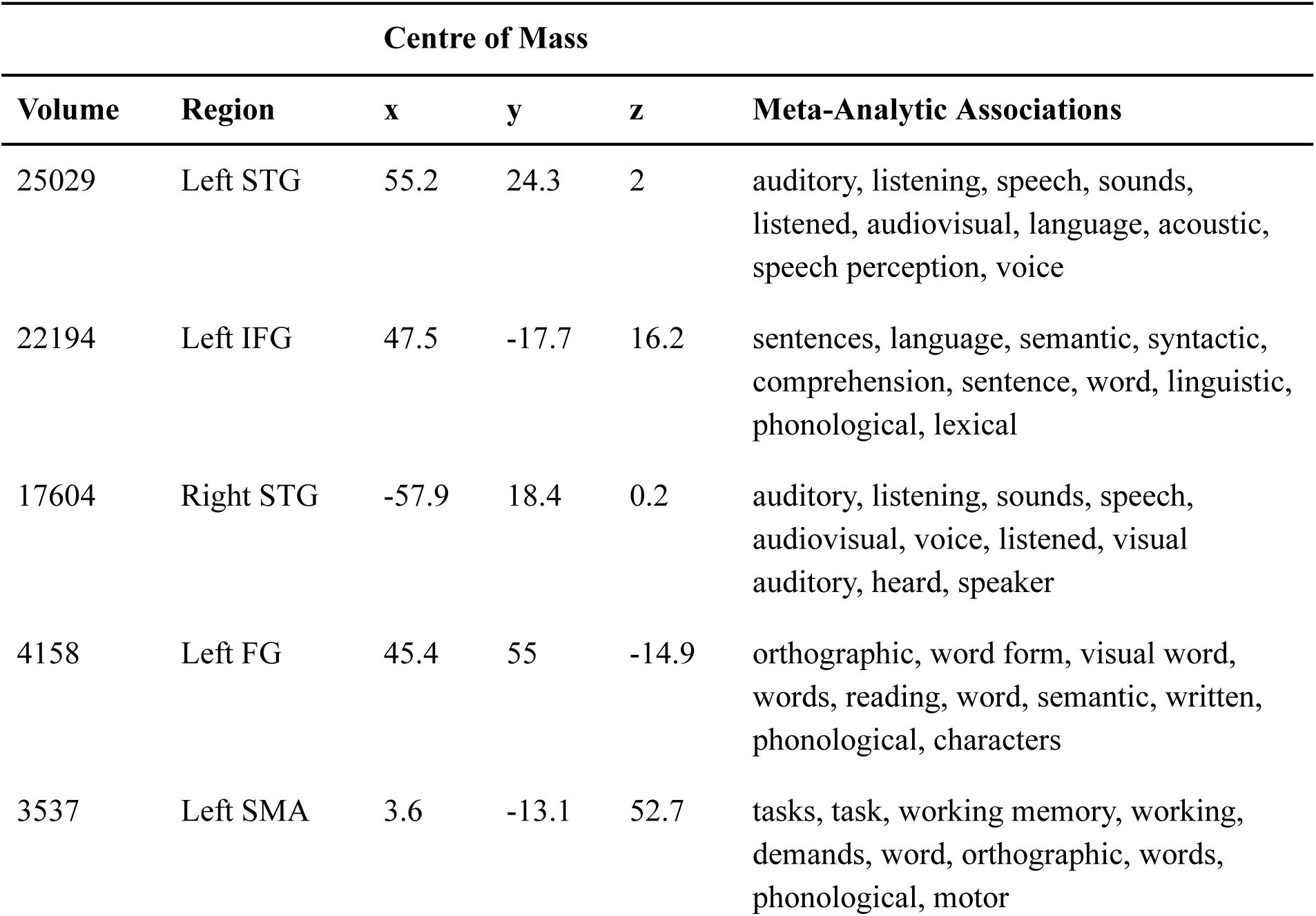
Clusters of significant activity from the neuroimaging meta-meta-analysis of language (Fig. 2, white outline). Significance was determined by a ‘group’ t-test using database as a covariate and thresholded at q ≤ 0.01 false discovery rate corrected with p ≤ 0.001 and a minimum cluster size of 50 voxels (400 µL). Volume is in microliters. Brain region abbreviations are the superior temporal gyrus (STG), inferior frontal gyrus (IFG), fusiform gyrus (FG), and supplementary motor area (SMA). The x/y/z centres of mass are in the Montreal Neurological Institute coordinate system. The meta-analytic associations are the top 10 functional terms at the given x/y/z coordinate from neurosynth.org (excluding, e.g., anatomical terms like ‘ifg’; all z-scores ≥ 6.74).

| Volume | Region | Centre of Mass |  |  | Meta-Analytic Associations |
| --- | --- | --- | --- | --- | --- |
|  |  | x | y | z |  |
| 25029 | Left STG | 55.2 | 24.3 | 2 | auditory, listening, speech, sounds, listened, audiovisual, language, acoustic, speech perception, voice |
| 22194 | Left IFG | 47.5 | -17.7 | 16.2 | sentences, language, semantic, syntactic, comprehension, sentence, word, linguistic, phonological, lexical |
| 17604 | Right STG | -57.9 | 18.4 | 0.2 | auditory, listening, sounds, speech, audiovisual, voice, listened, visual auditory, heard, speaker |
| 4158 | Left FG | 45.4 | 55 | -14.9 | orthographic, word form, visual word, words, reading, word, semantic, written, phonological, characters |
| 3537 | Left SMA | 3.6 | -13.1 | 52.7 | tasks, task, working memory, working, demands, word, orthographic, words, phonological, motor |

This apparent consistency raises the question as to why those specific anatomical locations support speech perception and language comprehension and why damage to them results in aphasia. One obvious answer is that they mostly involve regions adjacent to those in which acoustic information from the cochlea first arrives in the neocortex. It stands to reason that nearby regions would act upon that auditory information and transform it into the putative representations and processes that distinguish mere sound from language (like phonemes and phonology). It would be inefficient for distant regions to contribute because of the time delays and metabolic costs. Indeed, studies using ‘language localisers’ assert that ‘the language network’ processes only language information whereas more broadly distributed regions perform ‘extraneous’ and ‘autonomous’ functions ‘not necessary for linguistic processing’.^6,12–15^

### Averaging

An alternative model of the neurobiology of language explains differently the apparent anatomical consistency that is reflected by classical and contemporary models.^7,16^ It starts with the behavioural observation that ‘language’ is not only complex but also ambiguous at all levels, from speech sounds to semantics, syntax, and discourse. To resolve this ‘problem’, a large body of empirical evidence suggests that the brain makes use of contextual information stored internally (e.g., in the form of knowledge and expectations) and available externally (e.g., in the form of observable speech associated mouth movements and co-speech gestures;^17–20^ for further examples, see^16^) The memory and perceptual processes associated with internal and external context are distributed across the whole brain.^17–23^ These distributed networks appear to predict language input, e.g., in primary auditory cortex.^24^ Because context varies dynamically with each linguistic experience, language processing might never be the same twice.^16,25^ As such, the distributed regions involved in predicting language input would themselves be highly variable and dynamic.

Such a model is in the unenviable position of not conforming to Occam’s razor. That is, in addition to having to explain the consistency of regions appearing across studies, it must explain why we do not typically observe a whole-brain distribution of other regions engaged in language processing. One explanation is that they are concealed by existing methodological paradigms. For example, all neuroimaging studies rely on measures of central tendency, like averaging. In context-aware neurobiological models of language like that outlined in the prior paragraph, every word has distributed and variable activity patterns. Thus, averaging over different words, whether individually or in n-grams, sentences, or discourse, would reduce the activity in those peripheral regions with variable activity patterns to a low value.

This low value is even less likely to survive given the specific statistical analyses used. To illustrate, participants might listen to 100 different words during fMRI. The resulting data is a four-dimensional array of tens of thousands of ‘voxels’ collected at multiple time steps. Regressions are conducted in each of those voxels at the individual participant level, using a regressor that is a convolution of word onset times with a ‘canonical’ hemodynamic response function. Though this is not technically an average, it effectively acts as one by collapsing over different word representations. Resulting regression coefficients are then used to conduct group-level statistical analyses, again, at each voxel, resulting in average coefficients across participants. This is yet another level of collapsing over the different representations associated with each word as these may differ by participant. Finally, if any voxels exceed a statistical ‘threshold’ it is said that they are ‘activated’. However, thresholding requires prohibitive corrections for multiple comparisons because of the number of statistical tests that are done.^2^ Thus, connected peripheral voxels with legitimate but variable activity for any individual word are statistically unlikely to survive collapsing and averaging over word representations at the individual and group levels, particularly after thresholding. This would give the false impression that those vowels are not active and ‘extraneous’^3^ (Fig. 1).

**Fig. 1.**
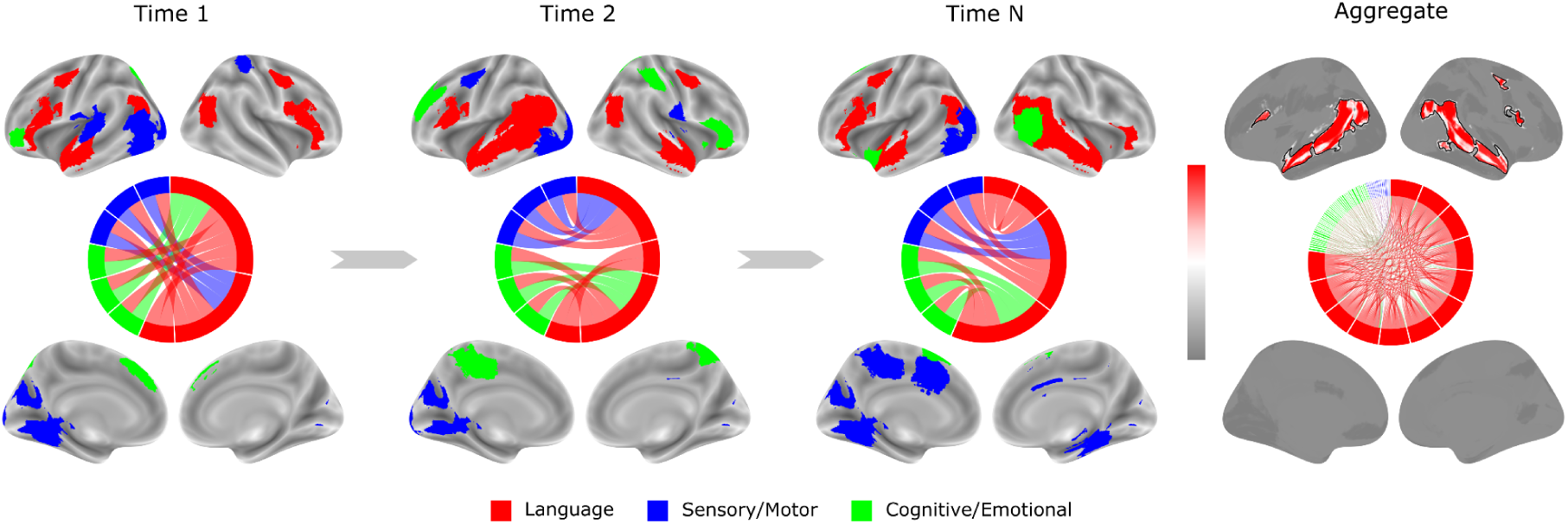
How averaging creates ‘language regions’ in neuroimaging studies. In this illustration, at Times 1–N, auditory cortices are consistently active as auditory input enters the neocortex here and is passed to nearby hubs (red brain regions). These hubs distribute information within themselves (red regions and ribbons in circle plots) but primarily to other regions (blue and green regions and ribbons). These peripheral connections vary across time more than auditory cortices. For instance, if sensory/motor (blue) and cognitive/emotional processes (green) are inseparable from word meaning or disambiguation, they would connect to the ‘language regions’—but only transiently. These connections shift with context and individual differences. Aggregating these data over time and applying thresholding preserves the consistently active auditory regions and hubs while averaging out more variable peripheral regions (rightmost panels). See Supplementary Materials for further details on how this illustration was generated.

**Fig. 2.**
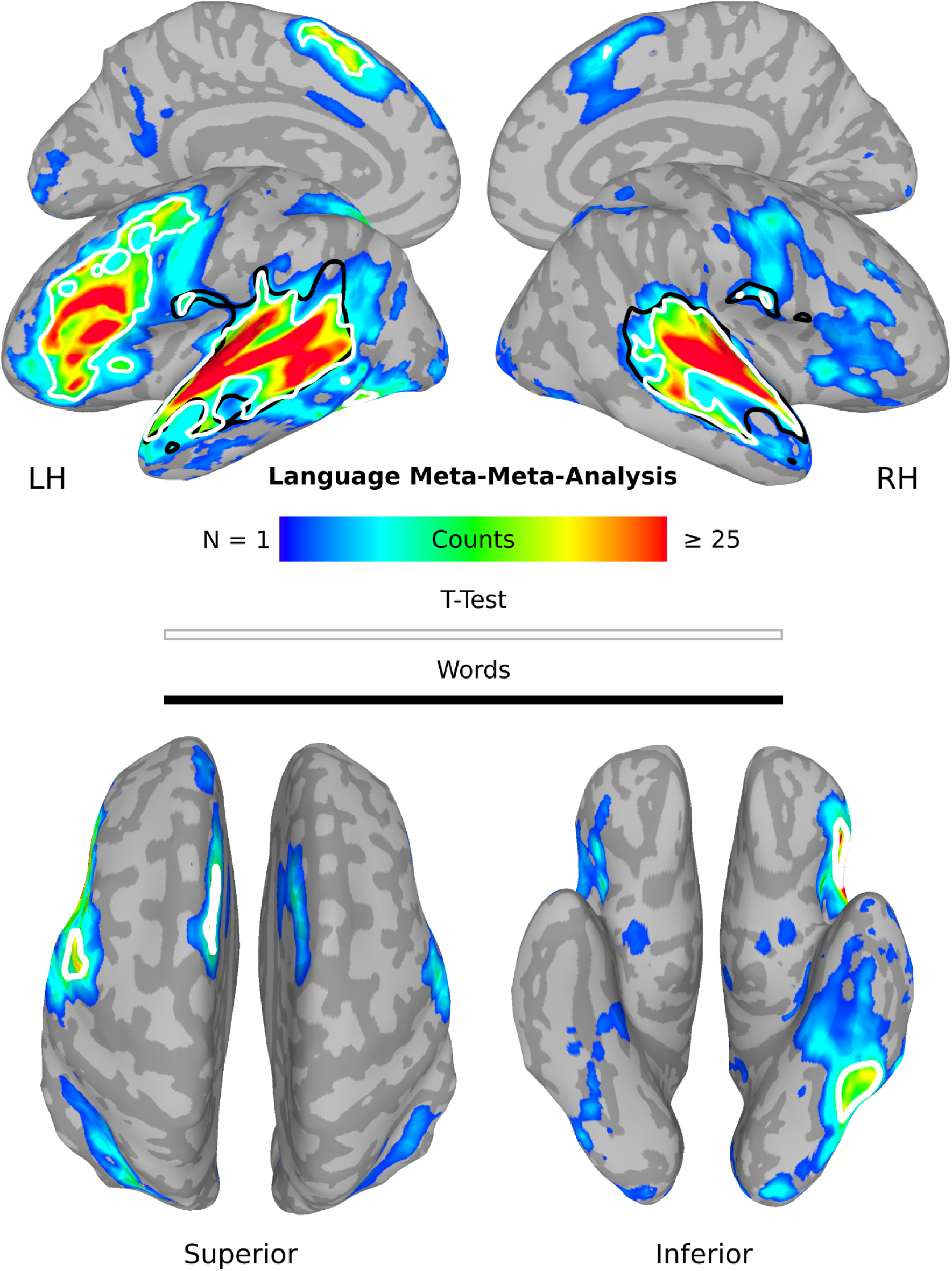
Neuroimaging meta-meta-analysis of language. Eighty-five meta-analyses of language representations (e.g., phonemes, words, sentences) and associated processes (e.g., speech, semantics, syntax) were conducted using the Neurosynth (N = 57) and BrainMap (N = 28) databases (Table 1). Each meta-analysis was thresholded at q ≤ 0.01 false discovery rate corrected for multiple comparisons using a minimum cluster size of 50 voxels (400 µL). This left 45 meta-analyses that each activated approximately the same regions. This can be gleaned from the number (N) of meta-analyses active in each voxel (colours) and a ‘group’ t-test false discovery rate corrected for multiple comparisons at q ≤ 0.01 with a minimum P value of 0.001 and a minimum cluster size of 50 voxels (400 µL; white outline; Table 2). The main effect of words from the movie-fMRI study is shown with a black outline for comparison (see Fig. 5).

### Distributed

Consistent with this picture, there is a large amount of evidence that a distributed context-aware model of the neurobiology of language is more consistent with reality when analyses do not indiscriminately average over heterogeneous word representations. To give an example, word processing can be considered to be instantiated in a distributed neural ensemble that incorporates experiences associated with word learning.^16,26^ Within this framework, verbs activate brain regions more associated with motion perception and limb movement whereas nouns activate regions more associated with object processing.^27^ This is also true of finer representations, whereby lexical processing that invokes auditory, visual, somatosensory, movement, and emotional related meanings activate brain regions partially associated with those processes, e.g., the transverse temporal gyrus, calcarine sulcus, postcentral gyrus, central sulcus, and insula.^28–30^ One cannot argue that these patterns simply represent post-linguistic ‘conceptual’ processing as they occur before this is possible, 50-150 ms after word onset.^31–35^

Results like these suggest that language processing is distributed across virtually the whole brain and that most of the neuroimaging evidence for this is obscured by averaging words from different semantic categories. A similar conclusion can be drawn from other linguistic representations. For example, overlearned language like ‘you know’ seems to not involve classical ‘language regions’ at all, explaining why they are often preserved after severe damage, as in global aphasia.^36,37^ A similar argument can be made not only for linguistic representations but other linguistic processes. For example, syntactic processing is arguably distributed,^38^ with different brain regions participating in disparate syntactic functions when these are not all lumped together.^39^ These include regions outside of the classical and contemporary ‘language regions’, e.g., the basal ganglia, pre-supplementary motor area, and insula.^40,41^

Compounding the issue, the act of averaging over all of these linguistic representations and processes typically conceals the distributed patterns associated with individual differences. For example, left-handed participants activate the right more than left motor cortices for verbs like ‘throw’, and the converse is true for right-handers.^42^ Similarly, professional hockey players activate premotor cortices for hockey-related sentences compared to fans and novices.^43,44^ More generally, we have known for more than 20 years that individual differences in patterns of language-associated activity do not correspond well to group averages.^45^ Group analysis by clustering shows extensive variability during language comprehension, with no one group of participants capturing the aggregate and individual participants varying on a spectrum involving the relative contribution of multiple neural structures, e.g., visual and sensorimotor regions.^46^

### Hubs

We return to the question of why ‘language regions’ or ‘the language network’ remain after averaging if standard methodological practices obscure most of the participating brain regions? One suggestion derives from the fact that the network organisation of the brain is neither random nor uniform. Rather, it has a small-world topology, characterised by the presence of hubs.^47^ Hubs are highly central regions (i.e., that have a high degree), with a large number of connections to other regions.^48^ This leads to the hypothesis that ‘language regions’ are a combination of auditory regions due to persistent audio input (at least in spoken language) and hubs necessary for the coordination of information processing in more dynamic, variable, and distributed peripheral regions.

Indeed, empirical evidence hints at the possibility that the regions identified in classical and contemporary models of the neurobiology of language are connectivity hubs. Structural MRI^49–51^ and ‘resting-state’ fMRI studies^52–62^ suggest that portions of superior and middle temporal and inferior frontal gyri are hubs. Though there are few studies measuring hubs using auditory or language stimuli, those that exist again suggest that these regions are functional hubs, encompassing even ‘early’ auditory cortices.^63,64^ However, across all structural, resting-state, and task-based studies, the location of hubs in this set of language-associated regions is quite variable. For example, hubs variously encompass portions of the anterior, middle, or posterior superior temporal gyrus and sulcus, suggesting they might be dynamic.^65^

### Hypotheses and Studies

To summarise, there is a conflict between the localizationist claims about ‘language regions’ and ‘the language network’ and data demonstrating that language processing occurs in a widely distributed set of brain regions. An alternative to localizationism is that these networks are not more obvious because averaging (and thresholding) over the more variable and distributed regions leaves only auditory regions and connectivity hubs that coordinate those more distributed peripheral regions. As we are not aware of any work empirically demonstrating this, we conducted four studies to test the following five hypotheses:

- **H1-Averaging**: Analyses that average over disparate linguistic representations and processes will show localisation to ‘language regions’.
- **H2-Distributed:** Analyses that do not average in this way will show specific linguistic representations and processes to be widely distributed throughout the whole brain.
- **H3-Hubs**: In analyses that average across linguistic representations and processes, the localised ‘language regions’ that survive averaging will be auditory processing regions and connectivity hubs.
- **H4-Dynamics**: Language-associated connectivity hubs will be shown to be dynamic, not fixed, and appear only in the aggregate.
- **H5-Connections**: Linguistic representations will be shown to be encoded primarily in the connections between ‘language region’ hubs and the brain periphery, rather than in individual hub or periphery regions.

### Meta-Analyses

For the first three hypotheses, we conduct a meta-analysis of neuroimaging meta-analyses, i.e., a ‘meta-meta-analysis’ of language.^66^ To test the first hypothesis, we attempt to demonstrate that studies that average over a variety of linguistic representations and processes consistently show activity in the same ‘language regions’ (H1-Averaging). However, this would not adjudicate between the veracity of the localizationist view versus the view that averaging conceals distributed regions. For this, we perform a second set of meta-analyses to determine if language processing in the brain is distributed when comparing the processing of gross linguistic representations, i.e., verbs and nouns (H2-Distributed; for other meta-analyses suggesting distributed processes, see^67,68^). Next, we conduct a novel meta-analytic centrality analysis to test the third hypothesis that the repeatedly activated ‘language regions’ expected in the meta-meta-analysis are hubs (H3-Hubs).

### Movie-fMRI

Even if the meta-meta-analysis reveals only ‘language regions’ and the verb and noun meta-analyses reveal a distributed pattern, the former cannot be derived by averaging over the latter. This and determining the relationship between ‘language regions’ and connectivity hubs must be demonstrated within participants. Furthermore, meta-analyses are typically static and cannot say anything about the dynamics of activity (H4-Dynamics), nor the representations carried by connections (H5-Connections). Thus, we used another approach, i.e., analysing movie-fMRI data from our ‘Naturalistic Neuroimaging Database’ or NNDb.^69^ First, we test the hypotheses that language processing during movie watching appears circumscribed to ‘language regions’ when using a measure of central tendency over words (H1-Averaging) but widely distributed when examining finer sensorimotor representations of those words (H2-Distributed). We tested the third hypothesis by determining the extent to which these averaged word regions are hubs, calculating this using multiple measures of voxel-wise network centrality (H3-Hubs). The fourth hypothesis that language-associated connectivity hubs are dynamic and not fixed was tested by making use of the temporally extended and context-rich nature of the movie-fMRI data (H4-Dynamics).

Specifically, the fourth hypothesis derives from the context-dependent nature of language use in the world (see ‘Averaging’ above). This view proposes that the forms of context used during naturalistic language processing vary dynamically and are coordinated by shifting spatiotemporal hubs. For example, speech-associated mouth movements may engage posterior superior temporal hubs, while iconic gestures recruit more anterior hubs.^17–20,24^ We hypothesised that coordinating hubs are not fixed, but vary over time, such that ‘language regions’ appear only collectively when aggregating across an extended processing period, but are not present at any particular time point (Fig. 1; H4-Dynamics). To test this, we derived voxel-wise network centrality from dynamic functional connectivity in overlapping windows, enabling analysis with and without aggregation over the fMRI data.

Positive results for H1–H4 would imply that language representations are distributed throughout the brain. However, an alternative explanation is that only information processing in ‘language regions’ is linguistic, while activity outside these regions is ‘non-linguistic’ and merely modulates the ‘language regions’. To adjudicate between these proposals, we tested the hypothesis that linguistic representations are encoded more strongly in the dynamic connections between language-related hubs and peripheral regions than in either the hubs or peripheral regions themselves (H5–Connections). This was assessed using novel voxel-wise encoding models that mapped contextual linguistic embeddings from GPT-2 onto fMRI-derived measures of local activation and functional connectivity.

Finally, though the meta-meta-analysis is immune from this criticism, it might be that the movie-fMRI results are somehow due to additional visual or other stimulation present in movies. For this reason, we attempted to replicate key results with audiobook-fMRI from the *Le Petit Prince* naturalistic fMRI corpus.^70^

### sEEG

Two questions that might arise are the extent to which these findings depend on the unique properties of fMRI, and whether brain activity outside putative ‘language regions’ truly encodes linguistic representations. Functional MRI is an indirect measure of neural activity, based on hemodynamics in 2–3 mm³ voxels, unfolding over ∼15 seconds and averaging across ∼500,000 neurons per voxel. This temporospatial blurring may make language appear more distributed than it is. To examine neural activity more directly, we collected data from 10 epilepsy patients implanted with electrodes while they listened to and watched audio stories, using stereoelectroencephalography (sEEG) to record local field potentials (LFPs). If nouns and verbs are similarly decodable in both ‘language’ and peripheral ‘non-language’ regions, this would support our second hypothesis (H2-Distributed). If LFPs in ‘language regions’ are more hub-like than not, meaning these regions have greater ability to encode both nouns and verbs instead of specializing for one word feature, this would support our third hypothesis (H3-Hubs), and together these results would lend further evidence for H5-Connections.

## Results

### Meta-Analyses

#### Averaging

We first conducted a neuroimaging ‘meta-meta-analysis’ to localise ‘language regions’ and to determine if putatively different language representations and processes activate similar ‘language regions’ across different meta-analyses. Such a result would be expected both under the hypothesis that ‘language regions’ are a self-contained localised system or network and the hypothesis that they are primarily a product of averaging. Specifically, 85 language-related meta-analyses were conducted (Table 1) and thresholded, correcting for multiple comparisons. Results were combined into a single brain image by count, followed by a one-sample ‘group-level’ statistical analysis, again correcting for multiple comparisons.

The results of this analysis revealed a circumscribed distribution of voxels activated in one or more language-related meta-analyses (Fig. 2; Table 2). On average, each voxel was activated in 6.36 of the meta-analyses, with a maximum single voxel activation in 45 meta-analyses (in the left dorsal superior temporal sulcus). Significant temporal lobe regions were activated across meta-analyses in both hemispheres, including the transverse temporal gyri, plana polaria and temporalia, superior temporal gyri and sulci, and posterior middle temporal gyri (Fig. 2, white outline; Table 2). Significant activity in the left hemisphere included the posterior inferior frontal gyrus, ventral and dorsal precentral sulcus and gyrus, and medial superior frontal gyrus (Fig. 2, white outline; Table 2).

#### Distributed

Second, we performed neuroimaging meta-analyses of ‘verbs’ and ‘nouns’ as an example to determine the spatial distribution of language representations when more specific linguistic representations are not averaged over, as is typically the case. This would provide evidence as to whether or not the ‘language regions’ in the meta-meta-analysis are likely to be the result of averaging. Indeed, verbs and nouns produce a whole brain distribution of activity (Fig. 3, colours) that is not encompassed by the significant ‘language regions’ from the meta-meta-analysis (Fig. 3, white outline). Specifically, significant activity for verbs included bilateral activity in the dorsal precentral sulcus and gyrus, central sulcus, postcentral gyrus, medial superior frontal gyrus, and medial temporal area (among other regions; Fig. 3, red). In contrast, nouns produced activity in bilateral occipital cortex, fusiform gyrus, and inferior occipital gyrus continuing into the inferior temporal gyrus (among other regions; Fig. 3, blue).

**Fig. 3.**
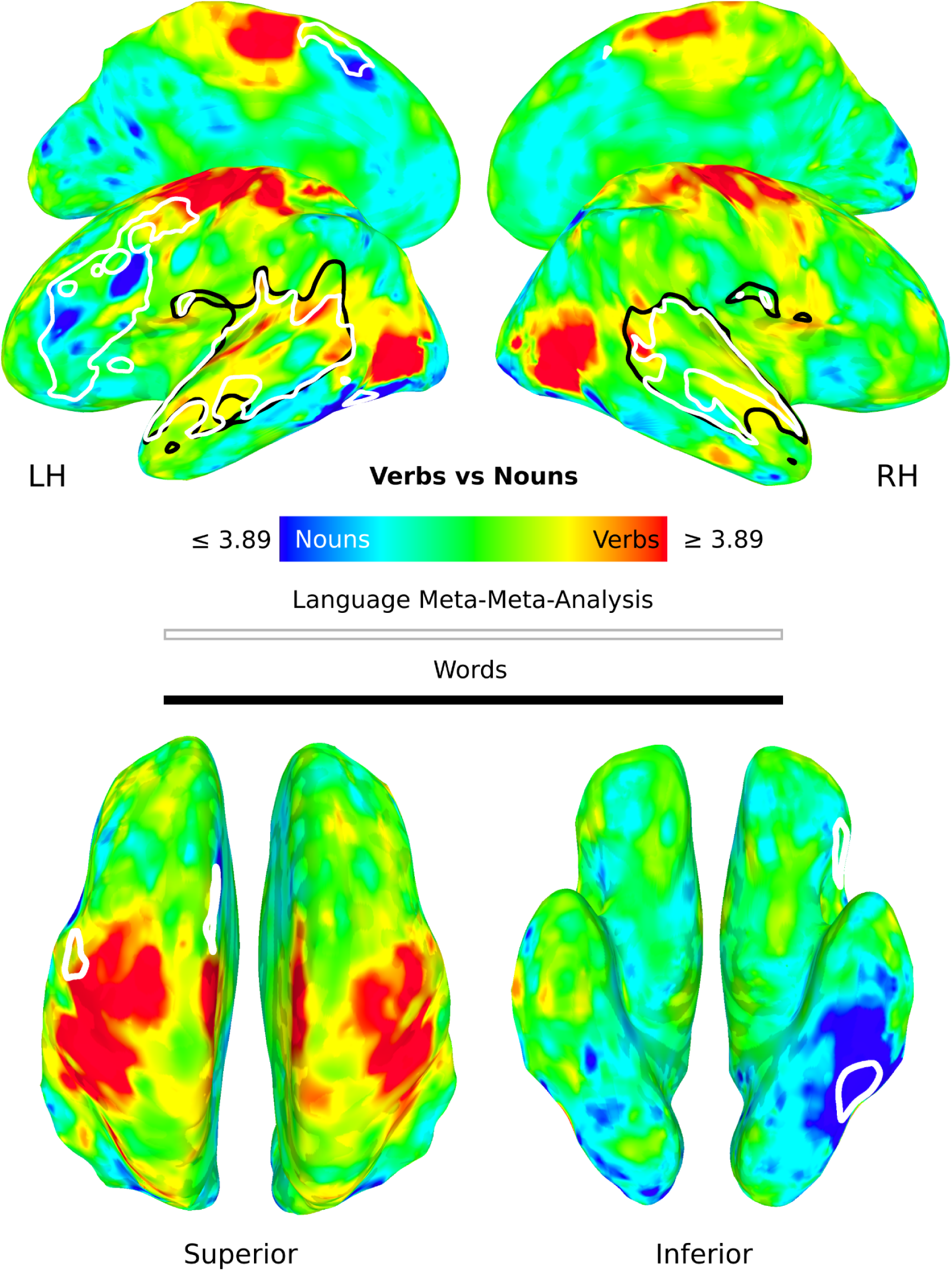
Neuroimaging meta-analysis of verbs and nouns. A meta-analysis of verbs (N = 662) and nouns (N = 889) was conducted using the NeuroQuery database. The z-scores were subtracted and presented unthresholded. Dark red and blue regions represent verbs and nouns, respectively, at a difference in the absolute value of the z-scores ≥ 3.89 or p ≤ 0.0001. Significant voxels from the language meta-meta-analysis in Fig. 2 are shown as a white outline. The main effect of words from the movie-fMRI study is shown with a black outline for comparison (see Fig. 5).

#### Hubs

Finally, we developed a new meta-analytic approach to quantifying degree centrality in order to evaluate whether or not the ‘language regions’ in the meta-meta-analysis are network connectivity hubs.^71,72^ If they are hubs, this would suggest that ‘language regions’ in the meta-meta-analysis remain after averaging over heterogeneous linguistic representations (like verbs and nouns) because they are hubs coordinating more distributed linguistic representations and processes. Specifically, we conducted 165,953 voxel-wise co-activation meta-analyses across 14,371 studies. As with functional connectivity, the regular co-activation of two or more regions suggests that those regions form functional connections or a network. Each meta-analysis was thresholded and combined by count (i.e., degree) as a measure of centrality and converted to a z-score for later thresholding.

The results indicate that a circumscribed set of regions in the task-driven brain have high centrality (Fig. 4). These include many regions in the superior and middle temporal lobes, posterior inferior frontal, and precentral regions from the meta-meta-analysis (Fig. 4, white outline). Indeed, the spatial correlation between the unthresholded centrality and language meta-meta-analysis was r = 0.52. When the centrality results were thresholded at z = 3.89 (p ≤ 0.0001), z = 3.29 (p ≤ 0.001), z = 2.58 (p ≤ 0.01), or z = 1.96 (p ≤ 0.05), they occupied 34%, 45%, 62%, and 76% of thresholded language meta-meta-analysis voxels, respectively.

**Fig. 4.**
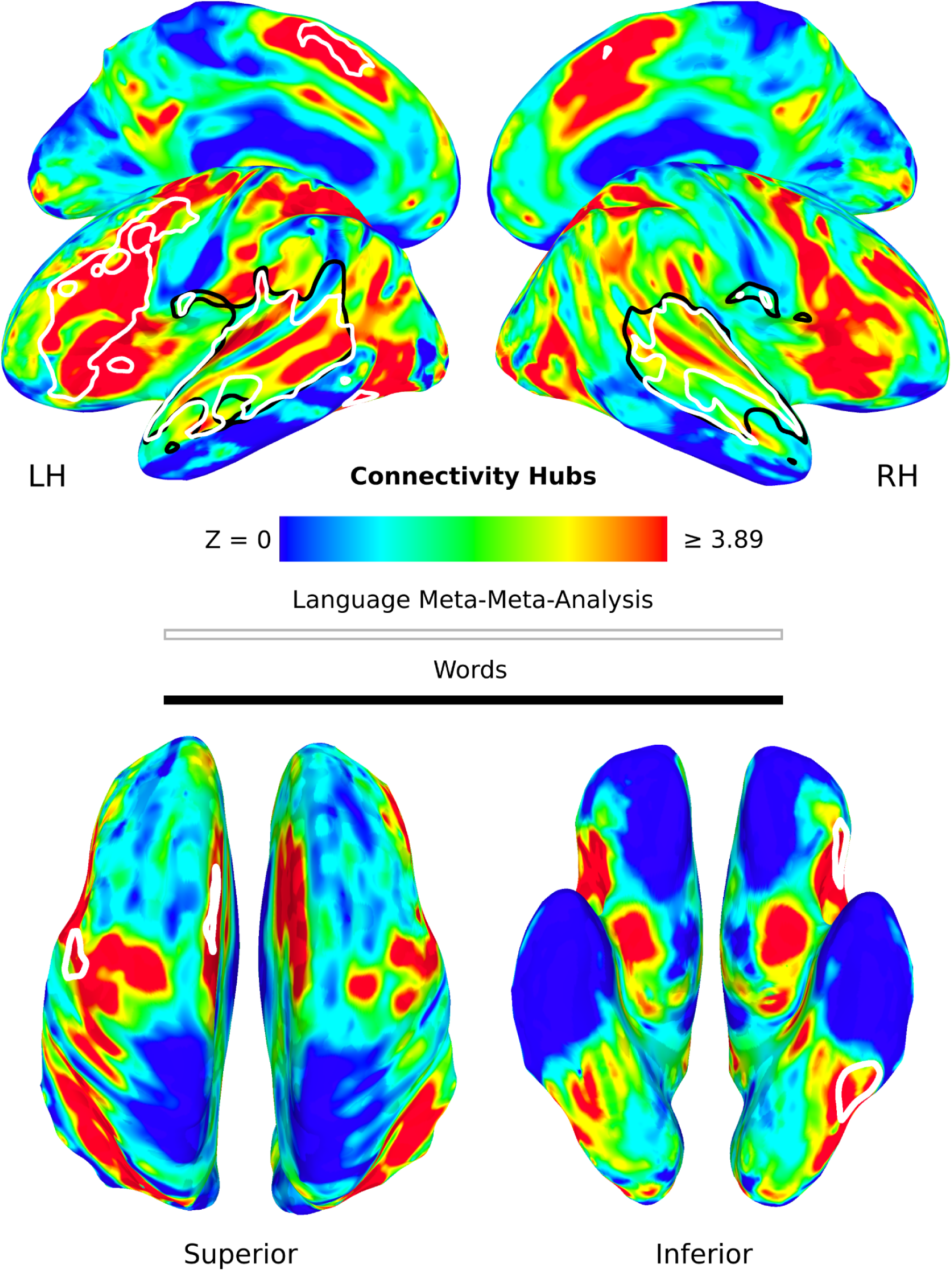
Neuroimaging meta-analytic connectivity hubs. Degree centrality was calculated using 165,953 voxel-wise co-activation meta-analyses across 14,371 studies, thresholded at q ≤ 0.01 false discovery rate corrected for multiple comparisons and combined by count. The mean and standard deviation across grey matter were used to convert counts to z-scores to make interpretation easier. Dark red regions are those that have z-scores ≥ 3.89 or p ≤ 0.0001. Significant voxels from the language meta-meta-analysis in Fig. 2 are shown as a white outline. The main effect of words from the movie-fMRI study is shown with a black outline for comparison (see Fig. 5).

#### Movie-fMRI

The results from the neuroimaging meta-analyses suggest that ‘language regions’ and ‘the language network’ might be the product of averaging over different word representations (Figs. 2-3), leaving only connectivity hubs (Fig. 4). However, the results are only suggestive because neither averaging nor connectivity hubs were generated within any one study, meaning that the observed findings might be due to some factor unrelated to our hypotheses.

#### Averaging

To overcome this issue and for other reasons discussed in the Introduction (see Hypotheses and Studies), we used movie-fMRI data from 86 participants to average over the heterogeneous properties of words in the brain, with the hypothesis that the results would be similar to the language meta-meta-analysis (Fig. 2). Specifically, we used duration and amplitude modulated regression at the individual participant level, modelling words and allowing them to be modulated by 11 sensorimotor experiential dimensions associated with the meanings of those words (i.e., auditory, foot-leg, gustatory, hand-arm, haptic, head, interoceptive, mouth, olfactory, torso, and visual dimensions).^73^ We also included sound energy, contrast luminance, and word frequency as nuisance modulators and regressors for words without sensorimotor ratings and nonword movie segments. For the group level analysis, the beta maps corresponding to the ‘main effect’ of words (henceforth ‘word’ or ‘words’) and the 11 sensorimotor modulators were entered into a linear mixed-effects model with beta, age, gender, and movie as fixed effects and participant as a random effect. The results were corrected for multiple comparisons using a cluster-size correction with multiple voxel-wise thresholds to α = 0.01 and a minimal cluster size of 20 voxels (540 µL).

Consistent with our hypothesis, word processing in the brain was spatially limited (Fig. 5). Regions of activity for words included the same bilateral superior and middle temporal lobe regions as in the meta-meta-analysis (Fig. 5, black outline). The predominant difference between these maps was the general lack of activity in the left inferior frontal and precentral gyri and sulci (compare the black and white outlines in Fig. 5). We included word frequency as a nuisance regressor to ensure that sensorimotor activity was not driven by this property (or ‘low-level’ auditory and visual features). However, it is well known that inferior frontal gyrus activity increases with decreasing word frequency,^74–76^ suggesting a role in selection and/or retrieval demands.^20,77–82^ To determine if the observed lack of activity was accounted for by word frequency, we conducted another amplitude-modulated regression and group linear mixed-effects model, this time without the 11 sensorimotor modulators. Directly contrasting sound energy with word frequency demonstrates that the latter nearly completely covers the left inferior frontal and precentral gyri and sulci, among other regions (and conversely provides evidence for the efficacy of the sound energy nuisance regressor; Supplementary Fig. S1).

**Fig. 5.**
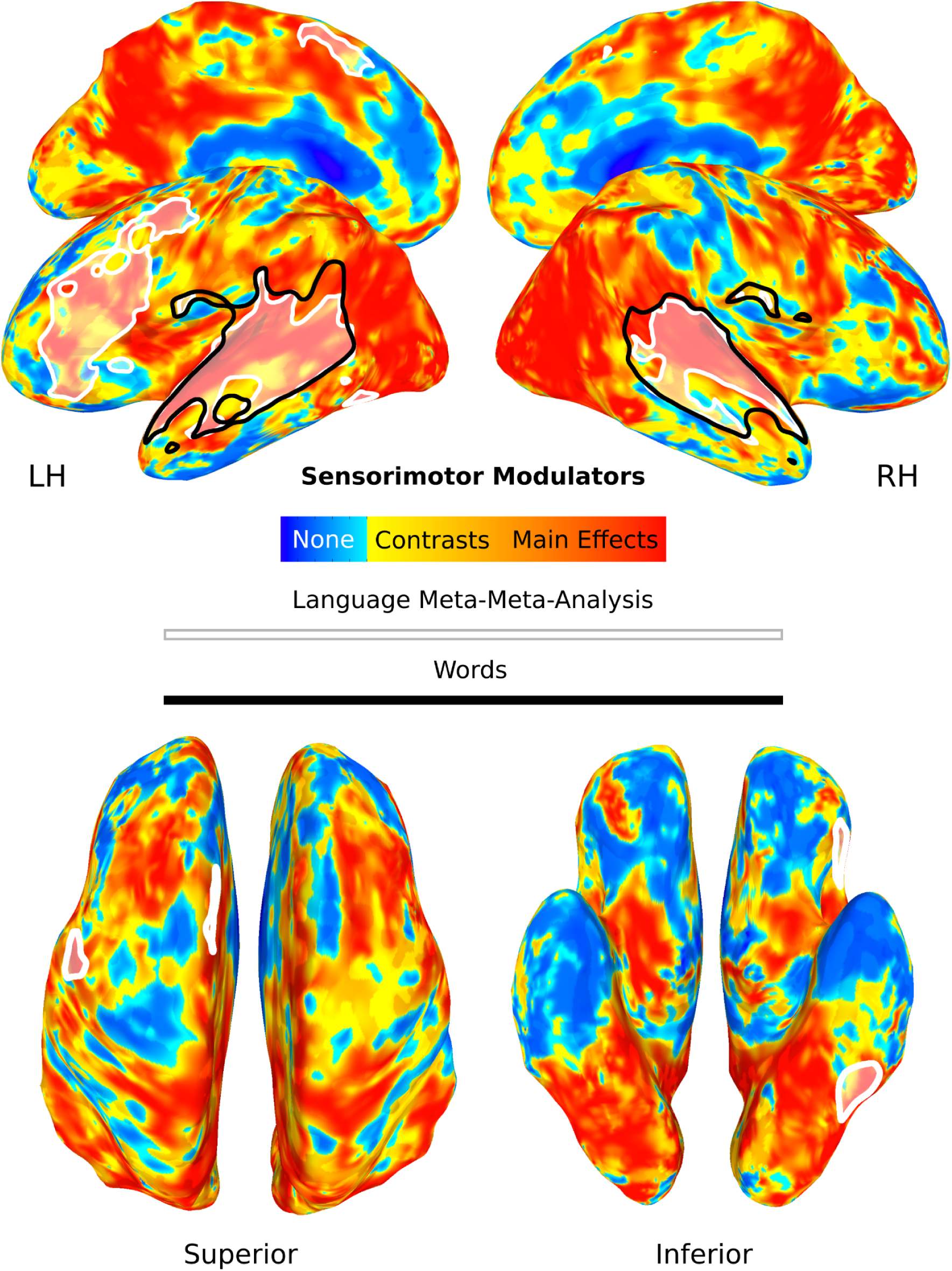
Distribution of activity for words and their sensorimotor experiential associations. The black outline corresponds to the main effect of words and the colours represent the 11 sensorimotor modulators of those same words. Specifically, the 11 sensorimotor modulators (e.g., foot/leg vs. baseline) are in red whereas the pairwise contrasts between modulators (e.g., foot/leg vs. hand/arm) are all shown in yellow. Insignificant voxels are in blue. All results are cluster-size corrected for multiple comparisons at α = 0.01 and displayed with a minimal cluster of 20 voxels (540 µL). The language meta-meta-analysis from Fig. 2 is included as a white shaded in outline for comparison to words.

For comparison, we next examined the similarity of activity patterns for word processing in this study with the results of the meta-meta-analyses. The percentage of thresholded word voxels that were also active in the language meta-meta-analysis was 52.64% (38,178 out of 72,522 µL; compare the black and white outlines in Fig. 5). If the voxels associated with word frequency are included (i.e., from the inferior frontal and precentral regions, see Supplementary Fig. S1), that percentage increases to 72.60% (52,650 of 72,522 µL). In contrast, the percentage of voxels from the 11 thresholded sensorimotor modulators of words (e.g., hand/arm) also in the language meta-meta-analysis was 7.42% (M = 5,380.36 of 72,522 µL, SD = 10,279.11; also see the discussion in the next section). The spatial pattern of activity for the unthresholded word map was correlated with the unthresholded language meta-meta-analysis at r = 0.51. In contrast, the 11 sensorimotor modulators of words did not correlate with the language meta-meta-analysis on average (with the mean r = -0.01, SD = 0.13).

#### Distributed

In parallel with the verb/noun meta-analyses (Fig. 3), we analysed the fine-scale sensorimotor properties of words within participants, evaluating the regional distribution in the brain. In this analysis, sensorimotor processing was distributed throughout much of the brain (Fig. 5, yellows and reds). Fig. 5 shows the main effect of the 11 sensorimotor modulators (e.g., foot/leg, Fig. 5, reds) and pairwise contrasts between modulators (e.g., foot/leg vs. hand/arm, Fig. 5, yellows) averaged and projected onto the brain, along with voxels that were not significant in any of these comparisons (Fig. 5, blues). In total, 36.47% of the brain was activated by sensorimotor modulators outside of voxels activated by words (478,386 of 1,311,687 µL), with each modulator activating an average of 4.32% of those voxels (56,653.37 of 1,311,687 µL, SD = 70,178.02). Similarly, when considering direct contrasts between modulators, 66.88% of the brain was activated outside of word voxels (877,203 of 1,311,687 µL), with each contrast activating an average of 4.31% of those voxels (56,584.14 of 1,311,687 µL, SD = 49,671.66). For comparison, words activated about 4.57% of the whole brain in total (62,856 of 1,374,543 µL). Finally, neither the spatial pattern of activity for the unthresholded sensorimotor (mean r = -0.02, SD = 0.24) nor the direct contrasts maps (mean r = -0.05, SD = 0.22) were correlated with the unthresholded word map on average.^4^

#### Hubs

In parallel with the overlap between the language meta-meta-analysis and the meta-analytic measure of centrality (Fig. 4), we expected regions associated with words to be hubs. To test this hypothesis, we constructed individual voxel-wise networks in just grey matter using a sliding-window approach and averaging over four measures of network centrality (degree, eigenvector, betweenness, and closeness). We clustered the centrality values using Ward’s minimum variance,^83^ which divided voxels in each time window into low- (given a value of one) and high-centrality clusters (given a value of two).

To examine hubs aggregated across time, we first averaged across all windows and performed a linear mixed effects model for the group-level analysis. The fixed effects were centrality (high and low), age, gender, and movie with participant as a random effect. A correction for multiple comparisons using a cluster-size correction with multiple voxel-wise thresholds was applied (again, using α = 0.01). Results show that high centrality voxels are significantly more active than low centrality voxels in most of the superior and middle temporal plane (Fig. 6, colours) and that the main effect of words overlap with these regions (Fig. 6, black outline). Indeed, 78.26% of the word voxels were high > low centrality (thresholded at α = 0.01 with a minimum individual voxel P value ≤ 0.001). Even when thresholding results further to include only the top 90% of the high centrality voxels, there was still a 39.00% overlap.

**Fig. 6.**
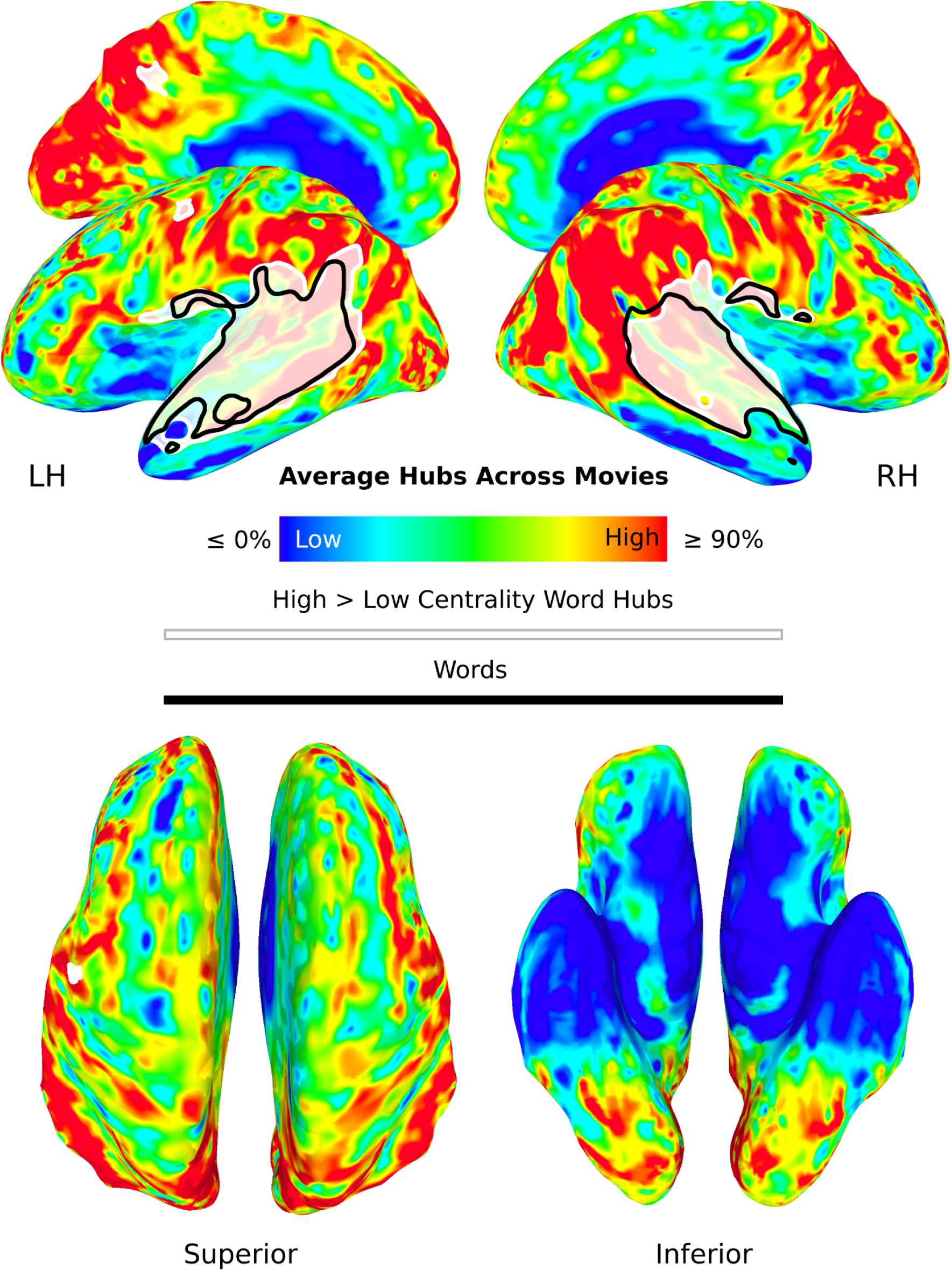
Distribution of high versus low centrality regions aggregated over time. Dynamic functional connectivity was conducted and colours show high minus low centrality regions averaged over time. Significant high > low regions that also exceed a 90% value to be considered hubs are in red. Spatial independent component and dual regression analyses were also conducted on the high and low centrality time series across participants to find sets of centrality hubs. Two components had a high correlation with the main effect of words (white shaded in outline) and were high > low centrality. The black outline corresponds to the main effect of words for comparison. All results were corrected for multiple comparisons.

The linear mixed effects model collapses over movies, not permitting the separation of high centrality hubs into possibly distinct sets. Thus, we next attempted to formally determine whether regions delineated by words formed a coherent set of hubs, independent of other sets in the aggregate across time. We accomplished this by performing group spatial independent component analysis (ICA) with 100 dimensions across participants’ dynamic high and low centrality time series. We then used dual regression to quantify the differences in high and low centrality, involving a direct contrast and thresholding using a correction for the 100 comparisons conducted. Finally, we computed the spatial correlation of each of the resulting contrasts with words, requiring a moderate or larger threshold (i.e., r ≥ 0.30). Results reveal two contrasts correlated with words (IC 8, r = 0.56 and IC 15, r = 0.53; two additional ICs survive at r ≥ 0.10, i.e., IC 5, r = 0.14 and IC 7, r = 0.25). These were both significantly more associated with high than low centrality and were combined for display purposes (Fig. 6, shaded white outline; see Supplementary Fig. S2 for examples of these and other ICs). Collectively, these two high centrality hubs shared 79.94% of their voxels with words and, together, were spatially correlated with words at r = 0.73.^5^

Thus, both the linear mixed effects and dual regression analyses suggest that voxels associated with words form a coherent set of connectivity hubs. However, we hypothesised that these only appear in the aggregate over time and do not exist as a whole on a moment-to-moment basis. That is, hubs are not fixed but are dynamic, e.g., moving around the superior and middle temporal lobes to coordinate distributed and varying peripheral regions. If this is the case, individual time windows should not be correlated with words as highly as in the ICA analysis. To examine this, we averaged across participants for each window for two of our movies separately, *500 Days of Summer* and *Citizenfour*. We thresholded the voxels in the resulting time series at 90% (i.e., ≥ 1.8) of the mean maximum value (i.e., 2) to isolate high centrality voxels. We then spatially correlated each thresholded time window with words. The mean correlation for *500 Days of Summer* was r = 0.03 (SD = 0.05) with only one time window having a correlation ≥ 0.30 (and only 10.55% of windows with r ≥ 0.10). The values for *Citizenfour* were comparable, with the mean correlation being r = 0.02 (SD = 0.04) with no time windows having a correlation ≥ 0.30 (and only 6.08% of windows with r ≥ 0.10). Performing ICA on these two movies produced similar results as those in the prior paragraph, with two networks correlated with words r ≥ 0.30 for each movie (IC 17, r = 0.58 and IC 35, r = 0.58 for *500 Days of Summer* and IC 31, r = 0.49 and IC 34, r = 0.66 for *Citizenfour*).

Collectively, these results suggest that the word hubs only appear in the aggregate over time (as in Fig. 6) and are present only in part on a window-by-window basis, despite nearly every moment of the movies containing language. To explore this further, we performed additional analyses to examine the connectivity between word hubs and the periphery at individual time windows. By the proposed model, when word voxels are acting as hubs, peripheral voxels should largely not be acting as hubs. To test this, we subtracted word voxels from the sensorimotor map (Fig. 5, black outline and colours), creating word and periphery masks. We computed the mean cluster assignment for each window for each participant across voxels in each of these masks, again considering a mean of 90% (i.e., ≥ 1.8) as high centrality. The word mask acted as a hub by this metric in 21.54% (SD = 9.66) of time windows on average across participants, lending support to analyses in the prior paragraph. In contrast, the peripheral mask had a value of ≥ 90% for only 0.07% (SD = 0.10) of time windows on average. When the word mask was a hub (with values ≥ 91.50% or 1.83), the peripheral mask was on average not a hub (with values ≤ 86.15% or 1.72). When the word mask was a hub, the average hub variance was 0.12 compared to 0.22 for the peripheral mask.

We performed the analyses described thus far on voxels labelled high and low centrality as we felt this was a conservative approach with regard to our hypotheses. That is, the more categorical approach is more likely to result in overlap and correlation with words at every time window (the null hypothesis). To understand if the results are comparable, we also conducted analyses on the averaged centrality maps directly thresholded at 90% to define high centrality regions across all 86 participants. Indeed, this showed something similar, i.e., on average 18.12% (SD = 12.20) of word mask voxels were high centrality in any given window. Described differently, ≥ 70.10% of windows had ≤ 10% high centrality voxels in the word mask and only 1.78% of the windows had ≥ 50% high centrality voxels. Yet, doing ICA on the 90% thresholded windows produced similar results as those above, this time with five networks correlated with words at r ≥ 0.30 (IC 1, r = 0.48; IC 3, r = 0.36; IC 4, r = 0.41; IC 12, r = 0.41; IC 23, r = 0.40; see Supplementary Fig. S3).

Finally, it might be the case that highly central word hubs are not connected to the sensorimotor periphery, e.g., if the periphery is ‘extraneous’ and processing is proceeding there independently somehow. To address this, we calculated the specific connectivity profiles of windows on an individual participant basis where the mean cluster assignment of the voxels from the word mask was ≥ 90% (i.e., ≥ 1.8, returning to the more categorical high/low windows) to the periphery defined, again, as the sensorimotor maps (i.e., the connectivity between the regions within the black outline and those in colour in Figs. 2-5). This analysis revealed that when the word mask was a hub, it shared an average of 43.99% (SD = 1.33) of connections with peripheral voxels across all participants. This helps explain the low spatial correlation between the thresholded windows and the word map and the relatively low percentage of word masks acting as hubs at any given time window above. That is, when word voxels are hubs at any given moment, they are connected to a periphery, driving down correlations with the word map that has been aggregated over time, thus excluding the less central and more variable periphery.

## Connections

The final analysis in the prior section suggests that word hubs are dynamically connected to a broad and distributed set of other brain regions. It is theoretically possible that the information processed in these peripheral regions is not language-related per se, but instead non-linguistic content that modulates activity in ‘language regions’ via connectivity. To address this, we adapted an approach from prior work aligning deep language models with brain responses to directly compare the predictive accuracy of local activation versus interregional functional connectivity.^84,85^ Voxel-wise encoding models were constructed to map contextual embeddings from GPT-2 layers 8 and 12 onto dynamic voxel-level activations in word hubs, peripheral activations, or functional connectivity between hubs and periphery, computed from moving time windows. Encoding used ridge regression with 5-fold cross-validation, and performance was assessed via mean Fisher-z correlations between predicted and actual fMRI signals in held-out data.

Group-level paired two-tailed t-tests on participants’ mean Fisher-z values revealed that voxel-wise prediction accuracy varied significantly across conditions (Fig. S4). Connectivity between hubs and periphery outperformed both hub and peripheral activations alone, with peripheral activations consistently yielding the lowest performance. Specifically, connectivity at layer 8 (mean Fisher-z = 0.22) was approximately 2.4 times more accurate than hub activations (mean = 0.09; t = 28.00, p < 0.001) and about 3.2 times more accurate than peripheral activations (mean = 0.069; t = 40.27, p < 0.001). Similarly, connectivity at layer 12 (mean Fisher-z = 0.12) was 3 times more accurate than hub activations (mean = 0.04; t = 33.83, p < 0.001) and 4 times more accurate than peripheral activations (mean = 0.03; t = 53.88, p < 0.001). These results suggest that peripheral regions are more than modulatory and that linguistic representations emerge from the connectivity between word hubs and peripheral regions, rather than from ‘language regions’.

### Audiobook-fMRI

Meta-analyses and movie-fMRI results support hypotheses that averaging conceals distributed patterns of language related activity, coordinated by connectivity hubs. Though the meta-analyses are immune from this criticism, it might be that the movie-fMRI results are due to visual stimulation. This is unlikely because nuisance regressors were used to control for the effect of extraneous auditory information and gross visual stimulation. Nonetheless, we attempted to address this concern by replicating results in an audio-only dataset comparable to the movie-fMRI dataset. Specifically, we used existing data collected at 3T from a group of participants (N=49) who listened to an audiobook (*Le Petit Prince*) for roughly the same amount of time as movie-fMRI movies (94 minutes).^70^

Key results replicated with approximately the same analyses, with the primary differences being that we did not need to include luminance nuisance regressors in the individual analyses or ‘movie’ as a covariate in the group analysis. Specifically, processing of all words was spatially limited to perisylvian regions whereas sensorimotor properties of words were distributed throughout most of the brain (Fig. S5). As with movie-fMRI, regions associated with averaged words are hubs. That is, high centrality voxels are significantly more active than low centrality voxels in the transverse temporal gyrus and superior temporal plane (Fig. S6). However, these hubs are dynamic and only appear in the aggregate over time. Dynamically, audiobook-fMRI hubs make connections with 38.24% of the rest of the brain. These connections carry the linguistic representations—not the hub or peripheral regions alone—with predicted connectivity yielding about 4–8 times greater accuracy than voxel-wise activations (Fig. S4; see the ‘Audiobook-fMRI’ section of the Supplementary Materials for detailed Methods and Results).

### sEEG

As noted, fMRI is an indirect measure of neuronal activity with a low temporal resolution. As such, it might inadvertently average over heterogeneous language processes and inadvertently make language appear more distributed. For these reasons, we used stereoelectroencephalography (sEEG) to record local field potentials (LFPs) from electrodes intracranially implanted in epilepsy patients listening to audio stories. We hypothesised that nouns and verbs would be similarly decodable everywhere in the brain, with ‘language regions’—defined as regions from a language meta-analysis—being more hub-like.

We focused on gamma-band activity, which has been shown to correlate closely with local firing rates.^86,87^ Gamma power, both within and outside putative perisylvian ‘language regions’ differentiated nouns from non-nouns and verbs from non-verbs (Fig. 7a–c). Encoding was stronger within ‘language regions’ (P < 0.001, Wilcoxon rank-sum test; Cliff’s delta = 0.86), but robust effects were also observed in regions outside those including, e.g., the anterior cingulate cortex, amygdala, hippocampus, insula, medial prefrontal cortex, orbitofrontal cortex, putamen, and thalamus (P < 0.001, permutation test). Based on receiver operating characteristic (ROC) analysis, a greater proportion of sites within language regions encoded both nouns and verbs compared to those outside (43% vs. 28%). Encoding strength for nouns and verbs was strongly correlated within language regions (r = 0.41) and outside them (r = 0.57; P < 0.001 for both; Fig. 7d), indicating that each electrode, encoding accuracy for nouns and verbs was relatively similar.

**Fig. 7.**
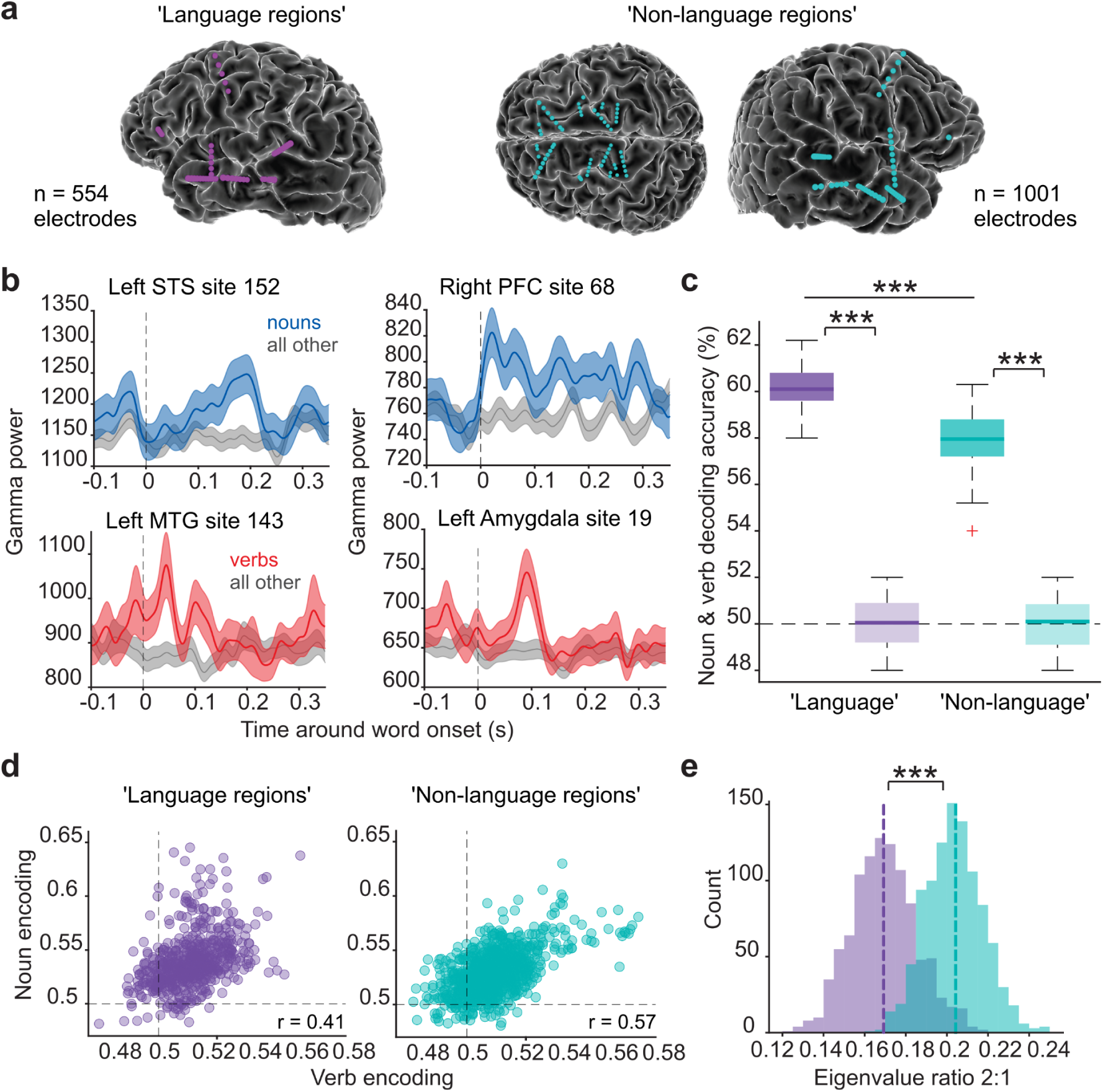
Brain-wide encoding of nouns and verbs from local field potentials. (a) Electrode locations from one example patient spanning both ‘language’ and ‘non-language’ regions. (b) Peri-event time histograms show gamma responses from four electrodes: superior temporal sulcus (STS) and middle temporal gyrus (MTG) ‘language regions’ and prefrontal cortex (PFC) and amygdala ‘non-language regions’. Lines represent word-averaged gamma power (± s.e.m.); dashed lines indicate word onset. (c) Decoding accuracy of nouns and verbs from gamma power was significantly above chance at both ‘language’ and ‘non-language’ sites (Wilcoxon rank-sum, P = 6.4 × 10^−26^); lighter boxes indicate shuffled-label controls (P = 0.0009, permutation test); chance = 50% (dashed line). (d) ROC AUCs for noun and verb decoding are shown for each electrode (dashed lines = chance); Pearson’s r = 0.41, P = 7.4 × 10^−24^ for ‘language’ and r = 0.57, P = 8.5 × 10^−87^ for ‘non-language’ regions. (e) Dashed lines show observed PC2/PC1 ratios from AUCs; bootstrapped distributions were generated from the dataset (see Methods). Wilcoxon rank-sum, P = 7.9 × 10^−263^.

The hub hypothesis predicts that contacts outside the ‘language regions’ should show a reduced tendency to have selectivity for both nouns and verbs, relative to contacts within language regions. One possible confounder is the intrinsic correlation between coding of nouns and verbs. To work around this confound, we examined the ratio of the second to the first principal component from neural and verb encoding accuracies for language and non-language regions (seen in Fig 7d); a larger ratio indicates a tendency to specialize in nouns or verbs, not both. (By contrast, differences in noise between regions would result in isotropic scaling). We find evidence for a shift in PC ratio. Specifically, the ratio of the second to the first PC is higher for contacts outside of the ‘language regions’ than for contacts inside (Fig. 7e, p< 0.001, Wilcoxon rank-sum test). Thus, the higher decoding performance for nouns and verbs that we observe in ‘language regions’ (Fig. 7c) is, in part, reflective of a more hub-like representation, since language regions are less selective for either nouns or verbs and more equally encode both compared to non-language regions (Fig. 7e).

## Discussion

### Summary

Classical and contemporary models of the neurobiology of language often suggest that there are a small number of fixed ‘language regions’.^5,6^ Yet, this is juxtaposed with significant evidence that language processing is distributed throughout the entire brain.^7^ To reconcile these differences, we tested the hypothesis that ‘language regions’ result from using measures of central tendency and thresholding across heterogeneous language representations and processes. Such averaging minimises the activation in regions with greater spatial variance while leaving activation in auditory input regions (in the case of vocal/heard languages) and connectivity hubs that coordinate those more variable peripheral regions (as illustrated in Fig. 1).

Indeed, using both neuroimaging meta-analyses and movie-fMRI, we show that language processing seems at first blush to occur in a very circumscribed set of fixed brain regions (Figs. 2 and 5). A meta-analysis of 85 individual meta-analyses of different language representations and processes shows that the same superior and middle temporal and posterior inferior frontal regions are repeatedly activated, with little spatial variance (Fig. 2). This same set of regions overlapped highly with those activated when 86 participants listened to words being spoken during one of 10 different movies while undergoing fMRI (Fig. 5, compare the black and white outlines; Supplementary Fig. S1).

This consistency in localisation disappears when words are no longer treated homogeneously (Figs. 3 and 5). That is, meta-analyses of even the gross distinction between verbs and nouns suggest that large swaths of brain tissue participate in language processing (Fig. 3). When making even finer distinctions in the movie-fMRI data, the distributed nature of processing becomes even more salient (Fig. 5, red and yellow). Specifically, individual (e.g., foot and leg, gustatory, interoceptive, and olfactory) sensorimotor properties of words produce distributed patterns of activity, averaging about 4% of the brain per each of 11 sensorimotor dimensions beyond word processing regions (over which those properties are averaged). Words consist of multiple sensorimotor properties and their conjoined activity encompasses up to 67% of the remainder of the brain. This includes regions important for processing action, emotions, interoception, movement, somatosensation, and vision (among others).

Thus, averaging and thresholding lead to misleading minimisation of activity in regions sometimes regarded as not language related or ‘extraneous’ (compare Figs. 2 and 3; Fig. 5). We hypothesised that this leaves only auditory processing regions centred around primary auditory cortex and regions of high connectivity that coordinate those distant but no longer observable regions. Indeed, most ‘language regions’ are connectivity hubs as defined by degree centrality in a large-scale meta-analytic connectivity analysis (Fig. 4) or the average of four different measures of centrality in the movie-fMRI data (Fig. 6; Figs. S2 and S3). However, additional analysis shows that these connectivity hubs are not fixed. Rather, they are spatiotemporally dynamic, with any individual hub occupying only some of what have become known as ‘language regions’ at any given moment and appearing together only when aggregating over time. On this moment-by-moment basis, language-related hubs are connected to a periphery of less central and more variable brain regions that are typically averaged away.

It could be the case that the interconnected distribution of hub and peripheral regions do not carry linguistic information. For example, could peripheral regions become engaged only by *post hoc* imagery of words and merely modulate activity in the putative ‘language network’? This does not seem viable, given that participants watching movies do not have time to imagine words. Indeed, we show that the connections between the dynamic hub and peripheral regions carry linguistic representations, not the individual hub or peripheral regions themselves (Fig. S4). Could such results be attributable primarily to the visual stimulation in movies? This is unlikely due to the analysis procedure that regressed out visual (and extraneous auditory) information, and the fact that the results fully replicate in an audiobook-fMRI dataset (Figs. S5 and S6). Lastly, could it be that the results are due to the poor temporal and spatial resolution of fMRI? This is contradicted by direct neuronal recordings of local field potentials, which show that brain regions thought to be specialized for language are actually hub-like, and that word categories can be decodable from gamma activity in both language and non-language regions, adding additional support that linguistic representations are distributed (Fig. 7).

### Local vs. Distributed

Collectively, these results beg for a reconsideration of localizationist accounts of the biological basis of language processing in the brain. Whether implicitly or overtly, it seems that the localizationist view of the 19th century, reconstituted in the imaging era of recent years, derives from confirmatory though potentially misleading results from thousands of neuroimaging studies. This is reflected in the widespread use of ‘language localisers’. These are ostensibly used for good reasons, to account for individual variation in the location of language regions,^88–90^ despite arguments against their usefulness.^91–93^ Regardless, a localiser task typically involves listening to intelligible language and to less intelligible language (among other variants), with analysis that requires averaging over many different kinds of linguistic representations and processes in both conditions, subtracting them, and thresholding. It is therefore not surprising, based on the arguments advanced here, that across 45 languages and even constructed languages like Klingon and Dothraki, localisers simply and repeatedly show the same regions as seen in Fig. 1 and 2.^94,95^ The unfortunate adoption of the localisionist view is also reflected in contemporary model building in which language processing is said to be supported by only a small number of fixed regions.^5,9,10,96,97^ The implication of our results is that these models are necessarily incomplete, in that they do not account for all regions or that they are wrong in claiming that only ‘language regions’ process language (and that language alone is processed by these regions).

There are several counterarguments that might be proffered in favour of maintaining more localizationist norms. Perhaps the more distributed results that we observe, particularly those in regions of the brain not considered core to language processing, are 1) due to ancillary processing like imagery that is not a core language function; or 2) limited to only ‘sensorimotor’ representations and ‘semantic’ or ‘conceptual’ processing. Though our data already counter these alternative explanations, we next provide additional argumentation, further solidifying that our distributed results are in fact due to representations and processes at the core of language processing. In the subsequent section, we further argue that a whole-brain model of the neurobiology of language is necessary for language comprehension and sketch a corresponding network architecture that might better account for the neurobiology of language than current models.

### Doing Language?

A sceptic might argue that the ‘language regions’ we observe do in fact only process language and that the other more distributed regions are doing something possibly independent and nonlinguistic, like post-perceptual imagery, conceptual processing, or thought. The methods and stimuli that we used in the movie-fMRI study when taken collectively help ensure that this is not the case. In the regression analysis, the sensorimotor properties were included as individual word amplitude modulators, meaning they are time-locked to word processing activity and thus not likely related to other processes. The model also included auditory and visual nuisance regressors to control for other features from the movies that might covary with those words (Supplementary Fig. S1). Moreover, speech in movies is continuous, without pauses between words, making it unlikely participants spent time imagining or conceptualising those words. Indeed, we show that *all linguistic representations* are carried by connections and not instantiated in individual regions. Finally, the results replicate in audio-only data and individual words can be decoded far afield ‘language regions’ from single neurons.

Further bolstering this argument, previous studies support the claim that those more distributed regions are doing something linguistic. Specifically, ‘language regions’ and other sensorimotor regions form networks during word processing and the latter are active within 50-150 ms after the word onset.^31–34^ Together, such results suggest that the sensorimotor component of the activation is part of and inseparable from the distributed representation of the words themselves.^26^ Our results support this view in that regions associated with words are connected to a large sensorimotor periphery on a moment-to-moment basis. This view is also consistent with results in other domains like vision and memory showing that representations are not confined to but are maintained across reciprocally interconnected neurons and regions.^98–100^

In contrast, the strongest claims involving ‘language localisers’ suggest that the putatively domain-general ‘multiple demand network’ (MDN) ‘shows no sensitivity to linguistic variables’.^101^ This argument creates a false and sterile dichotomy.^102^ First, the MDN is often sensitive to ‘linguistic information’ albeit at reduced levels.^12,13,101,103–106^ Second, analyses are done by averaging over large regions with different connectivity and cytoarchitecture profiles,^107^ comprising about 25% of the grey matter voxels in the brain.^6^ This would lead to less sensitivity for detecting ‘linguistic information’, particularly given the demonstrations herein, e.g., that this activity is dynamic and variable. Finally, MDN regions are appropriately named as they have some of the highest measures of functional diversity and ‘neural reuse’ in the human brain.^102,108–112^ These functions are most certainly not limited to domain-general processes. For example, premotor cortices play clear roles in speech perception and language comprehension.^113^ Indeed, as we expand on in the next section, these putatively non-language regions do contribute specifically to language processing when analyses do not average over individual language representations and processes.

### More Than Sensorimotor?

It is also unlikely that our argument that ‘language regions’ are the results of averaging is specific to only ‘sensorimotor’ representations and ‘semantic’ or ‘conceptual’ processing. That is, the entire set of distributed regions beyond the putative ‘language regions’ has been variously demonstrated to be involved in many other language representations and processes when they are probed. Indeed, our encoding analysis covers linguistic representations other than word semantics.^85,114^ To give further examples, people with global aphasia who are missing the entire ‘language network’ can still produce some formulaic language (or multiword expressions), suggesting these linguistic representations are instantiated outside of ‘language regions’.^37^ Despite this, there are fewer than five neuroimaging studies that we are aware of that distinguish formulaic language as a different type of linguistic representation. Furthermore, in a recent study,^36^ we have shown that novel sentences are processed in ‘language regions’ (when averaged over) but when these same sentences are overlearned over 15 days, becoming more formulaic, they are processed outside of these regions (primarily in the central sulcus and subcortical structures).

Many other examples can be provided that non-‘language regions’ are involved in language beyond their role in sensorimotor representations. To give a non-exhaustive overview, orbitofrontal cortex, among other regions, plays a role in indirect speech act comprehension.^115^ The dorsolateral prefrontal cortex (part of the putative MDN) plays numerous specific roles in ‘discourse management, integration of prosody, interpretation of nonliteral meanings, inference making, ambiguity resolution, and error repair’.^116^ Dorsal medial prefrontal regions are involved in understanding speech acts, nonliteral meanings, and emotionally and socially laden language.^117^ These regions and the precuneus are involved in various aspects of generating and updating language-based situation models.^118,119^ More generally, these regions comprise aspects of the ‘default mode network’ which have been directly linked to ‘language regions’, inner speech, and language comprehension.^120–124^

Other regions, like the anterior cingulate and parietal cortices, help implement necessary adaptive language control processes.^125^ Entire distributed motor systems play many roles in supporting speech perception.^113^ The insula and limbic structures like the amygdala are involved in affective prosody processing.^126–128^ Other subcortical structures like the basal ganglia,^40,129^ thalamus,^130–132^ and cerebellum^133^ play numerous linguistic roles, including the processing of speech, semantics, and syntax. Visual regions like the putative ‘visual word form area’^134^ and face and motion processing regions^135,136^ play roles in audio-only speech perception, even when there is no visual information available. Indeed, it has long been recognized that there is a ‘basal temporal language area’ in the fusiform gyri in occipital cortices associated with receptive aphasia in the auditory modality.^137–142^

### NOLB Model

Claiming that these various distributed language representations and processes are somehow non- or extra-linguistic or, worse, ‘merely’ pragmatic is problematic. It threatens to reduce our real-world understanding of language to some core elements that in themselves cannot explain how the human brain manages to generate and extract information through language, e.g., to syntactic recursion^143,144^ or semantic compositionality.^145^ However, we suggest there is an even more fundamental reason for rejecting linguistic and nonlinguistic dichotomies for a whole-brain neurobiological model. That is, doing so is part of an account that helps resolve a fundamental problem in speech perception and language comprehension, namely, how the brain overcomes linguistic ambiguity. Next, we elaborate on a mechanistic model (that we sketched out in the Introduction) by which the brain uses both linguistic and nonlinguistic contextual information to predict language representations, constraining interpretation.^16^

Language representations and their associated processes are ambiguous at all levels (for a review, see).^16^ For example, there are no known acoustic features that unambiguously distinguish phonemes (called the ‘lack of invariance problem’). Words are not only homonymous (e.g., ‘bat’ and ‘bank’) but most are polysemous (e.g., the word ‘set’ has more than 450 meanings). Similarly, sentences and discourse can be syntactically and/or semantically ambiguous. How does the brain resolve all these ambiguities? We and others have taken proposals from Helmholtz in vision (‘unconscious inference’, circa the 1860s) and Stevens and Halle in speech (‘analysis-by-synthesis’, circa the 1960s) and proposed that language ambiguity necessitates, in more modern parlance, prediction.^16,113,146^

Specifically, the brain uses the ubiquitous amount of internal context (in the form of memories) and external or observable context it has available in the real world to make predictions about forthcoming language representations. To give an example, observable speech-associated mouth movements precede associated auditory information by about 100-300 ms.^147^ These can be used to make predictions about the acoustic patterns subsequently arriving in primary and surrounding auditory cortices and constrain the interpretation of which phonemes are intended. We have shown that this is how audiovisual speech perception works, involving feedforward and feedback network interactions between visual, ventral pre- and primary motor cortices, and posterior superior temporal cortices (among others).^17,19^

Because language is ambiguous and requires continual predictions derived from context, and context is always variable, the brain regions involved in speech perception and language comprehension will necessarily be highly variable. As these distributed regions are making predictions, they form connected networks. The only common denominator in these spatially variable and distributed networks are auditory processing regions and connectivity hubs coordinating them. This has two implications relevant here: First, because, e.g., visual and motor regions are predicting intended phonemes, the distinction between ‘linguistic’ and ‘nonlinguistic’ is fuzzy at best. Second, averaging over the various representations and processes occurring in different and more variable distributed networks leaves only the aforementioned auditory and hub regions after thresholding.

What kind of network architecture can accommodate such a model? A classically modular architecture is not capable of achieving the flexibility needed to accommodate continually changing contexts and associated networks. An alternative architecture incorporates core-periphery structures that combine two network dynamics: The core is a collection of stable hubs that control a set of flexible regions, or periphery, that manifest remarkable temporal variability.^148,149^ Core-periphery networks allow for high complexity, robustness to perturbations and rewiring of connections, and recovery following injury.^149,150^ Our data appear to fit into a core-periphery framework with the identified ‘language regions’ corresponding to dynamic cores and the rest of the distributed activity (that is typically averaged away) constituting a dynamic periphery. It is left to future work to test this architecture with a whole-brain, voxel-wise core-periphery algorithm.^151^

### Implications

The studies and results here suggest that the historical neo-localizationist view of the neurobiology of language needs radical revision.^16,152^ By obscuring the distributed and whole-brain nature of language processing systems, static neuroimaging studies have inadvertently reinforced inferences made from lesion analyses in the 19th century. The latter suggest that gross language dysfunctions are caused by damage to a small set of fixed regions. Similarly, neuroimaging analyses based on averaging over linguistic ‘apples and oranges’ and thresholding, give the false impression that language processing occurs in a small set of fixed regions. Results here suggest that both the language problems caused by injury and the localisationist inferences that can be drawn from neuroimaging can be explained by considering that ‘language regions’ and ‘the language network’ are some combination of auditory processing regions and connectivity hubs. This implies that aphasia is the result of damaging the core regions and their connections that coordinate a whole-brain distribution of regions that process language and not the result of damaging ‘language regions’. Indeed, empirical evidence demonstrates that damage to connectivity hubs is more likely to underlie various disorders, including aphasia.^153–158^ This implies that a different (perhaps core-periphery) model of the neurobiology of language is needed to replace contemporary models to help advance treatment for aphasia (where effect sizes are low on average).^159^ This might involve focusing therapy around individualised and preserved linguistic representations and processes in the periphery.

More generally, it is perhaps obvious that our conclusions about averaging and thresholding can be extended to every domain in which psychological ontologies are probed with neuroimaging data.^46,160^ It is hard to think of any neuroimaging study in which stimuli do not contain multiple representations or processes that are mathematically integrated before thresholding. Therefore, what is revealed by most studies and neuroimaging meta-analyses is more likely to reflect underlying connectivity hubs rather than the full distribution of regions involved. This conclusion probably scales with the complexity of the behaviour under investigation. For example, visual object processing studies average over many different stimuli whose sensorimotor properties and affordances greatly vary, leaving only core regions involved in complex visual processing (like the fusiform gyrus). In contrast, language and social processing are at a pinnacle of human functioning and likely require more dynamic hubs orchestrating far more regions given the complexity of these processes. This implies that we need much more consideration of the representations and processes engaged by the stimuli and tasks we give participants and considerably more emphasis on individual differences.^160–163^

## Methods

### Illustration

Fig. 1 is the result of an informal simulation intended to illustrate an alternative explanation for why specific regions might consistently appear in standard neuroimaging studies involving language-related contrasts—not because they are the only ‘language regions’ in the brain, but due to the effects of averaging. It is ‘informal’ because it is largely qualitative in nature and was conducted primarily to generate images using standard neuroimaging software, which would be difficult to reproduce manually. A more comprehensive study would be required to complete a more quantitative simulation. Nonetheless, considered as a set of results, the simulation is consistent with the primary hypotheses outlined in the Introduction.

To generate Fig. 1, we randomly selected nine of 12 parcels from a publicly available probabilistic map of a language localizer task involving 220 participants (Fig. 1, red). We then randomly chose 12 additional regions outside this putative ‘language network’ from a 200-region parcellation atlas,^164^ grouping them into two sets of six. Random connection weights were assigned between the language localizer regions and each set. These spatial and connectivity patterns represent ‘language’ hubs linked to sensory/motor and cognitive/emotional peripheries (Fig. 1, blue and green). The peripheries and their labels are arbitrary and included for illustrative purposes only—they could have involved any set of brain regions. We simulated 10,000 time points (Fig. 1, left three columns), aggregated the data, and thresholded the result at the 90th percentile to retain the most consistently active voxels (Fig. 1, rightmost column). More details are provided in the ‘Illustration’ section of the Supplementary Materials.

### Meta-Analyses

#### Averaging

To conduct the neuroimaging ‘meta-meta-analysis’, we manually searched the Neurosynth^165^ and BrainMap^166^ databases (accessed in August 2022) for available terms and studies related to language representations (e.g., ‘phonological’, ‘words’, ‘sentences’) and associated processes (e.g., ‘speech’, ‘semantics’, ‘syntax’). We excluded terms and study descriptions pertaining to music and reading or having an obvious visual element. This resulted in 57 Neurosynth terms and 28 BrainMap searches that were used to assemble the meta-meta-analysis (see Table 1).

Specifically, the Neurosynth meta-analyses are already generated and we simply downloaded them from neurosynth.org in Montreal Neurological Institute (MNI) stereotaxic space^167,168^ with a false discovery rate corrected threshold of q ≤ 0.01 (from dataset version 0.7, released July 2018).^165^ These had been constructed by extracting 507,891 activation coordinates reported in 14,371 neuroimaging articles and collating a list of 1,334 high-frequency terms that occur in more than 20 of those articles. Coordinates for each high-frequency language-related term are aggregated and compared to the coordinates reported for articles without those terms (Table 1, left). For BrainMap meta-analyses, meta-data from the ‘Sleuth’ application (version 3.0.4) was used to search for papers (Table 1, right) and corresponding coordinates, and the ‘GingerALE’ application (version 3.0.2) was used to perform activation likelihood meta-analyses on those coordinates.^166,169–171^ These were done in MNI space and thresholded using false discovery rate correction of q ≤ 0.01 to match meta-analyses done in Neurosynth.

Each of the resulting 85 meta-analyses was further preprocessed to have a minimum cluster size of 50 voxels (400 µL). The spatial pattern of activity for the combined Neurosynth meta-analyses was highly correlated with the BrainMap meta-analyses (r = 0.76). Thus, the results were combined into one neuroimage by count using ‘3dmerge’ available in the AFNI software package.^172^ Additionally, we conducted a one-sample ‘group level’ statistical analysis with ‘software package’ as a covariate (i.e., Neurosynth or BrainMap) and a minimum of five contributing meta-analyses using ‘3dttest++’, also available in AFNI.^172^ Results were thresholded using a false discovery rate correction of q ≤ 0.01 and a minimum cluster size of 50 voxels (400 µL). We used the online database at neurosynth.org to provide functional descriptions of the resulting clusters of activation (from dataset version 0.7, released July 2018).^165^ Specifically, we give the top 10 significant functional terms at the centre of mass coordinates, excluding anatomical terms like ‘heschl gyrus’, ‘inferior’, and ‘superior temporal’.

#### Distributed

To determine the extent of language processing in the brain when not averaging over all representations, the NeuroQuery database was queried for ‘verbs’ and ‘nouns’ as an example of differentiable representations that are often averaged over, e.g., as would necessarily be the case in the ‘sentences’ meta-analysis (accessed, August 2022).^173^ Note that we chose verbs and nouns primarily because few other specific linguistic contrasts are available for meta-analysis, and because this choice allowed us to perform a parallel sensorimotor analysis with our movie-fMRI data. Unlike Neurosynth, NeuroQuery produces meta-analyses that are predictions of where in the brain a study about the input terms is likely to report activity. We used this approach over Neurosynth because it can make use of coordinates from nearly 10 times more articles which would presumably make results more robust for relatively less frequently studied topics. In particular, NeuroQuery produced a meta-analysis from 662 articles for the term ‘verbs’ (compared to 107 in Neurosynth) and 889 for ‘nouns’ (compared to 100 in Neurosynth). Results are presented unthresholded in Fig. 3. To determine significant regions of activations reported in the Results section, the meta-analyses were subtracted from each other and thresholded at z = 3.89 (p ≤ 0.0001), excluding any overlap and using a minimum cluster size of 50 vowels (400 µL). The patterns of activity for these thresholded results were similar to but more robust than the q ≤ 0.01 false discovery rate correct ‘verbs’ and ‘nouns’ meta-analyses from Neurosynth.

#### Hubs

We developed a new meta-analytic network connectivity hub metric to examine whether the resulting ‘language regions’ from the meta-meta-analysis are hubs.^71,72^ First, we used the Neurosynth software package to conduct 165,953 voxel-wise co-activation meta-analyses across 14,371 studies. Similar to the logic of functional connectivity, the regular co-activation of two or more regions is suggestive that those regions form functional connections or a network. Each meta-analysis was thresholded at q ≤ 0.01 false discovery rate corrected and combined by count using ‘3dmerge’ available in the AFNI software package.^172^ This corresponds to the number of connections (edges) that a voxel (node) has and, thus, is a meta-analytic equivalent of degree centrality. The resulting map was spatially converted to z-scores using a white matter mask generated by Freesurfer to limit results to grey matter.^174,175^ This is presented unthresholded in Fig. 4. Hubs were defined as voxels with 3.89 standard deviations more connections than the mean (equivalent to p ≤ 0.0001). Overlaps with meta-meta-analysis results were tested at this z-value as well as z = 3.29 (p ≤ 0.001), z = 2.58 (p ≤ 0.01), and z = 1.96 (p ≤ 0.05).

### Movie-fMRI

#### Neuroimaging

Our fMRI data was derived from 86 participants who watched one of 10 previously unseen full-length movies from 10 different genres (42 females, age range of 18–58 years, M = 26.81, SD = 10.09 years). All participants were right-handed, native English speakers with normal hearing and vision (corrected), without claustrophobia or neuro-psychiatric illness, and not taking medication, all determined by self-report. The study was approved by the ethics committee of University College London and participants provided written informed consent to take part in the study and share their anonymised data.

Functional MRI scans were collected on a 1.5T Siemens MAGNETOM Avanto using a 32-channel head coil and a multiband EPI sequence (TR = one second, TE = 54.8 ms, flip angle of 75°, 40 interleaved slices, resolution = 3.2 mm isotropic, multiband factor = four, no in-plane acceleration), with breaks determined by participants about every 40 minutes. The movies were one hour and 56 minutes long on average, meaning there was a mean of 6,998.20 TRs (SD = 1068.92 TRs; range is 5,470 to 8,882 TRs). A 10-minute T1-weighted anatomical scan was collected after the functional scans using an MPRAGE sequence (TR = 2.73 seconds, TE = 3.57 ms, 176 sagittal slices, resolution = 1.0 mm).

Unless otherwise noted, all preprocessing and data visualisation was done with the AFNI software package and specific programs are indicated where appropriate^1727^. First, anatomical images were corrected for intensity non-uniformity, deskulled, and nonlinearly aligned to an MNI template (i.e., MNI152_2009_template_SSW.nii.gz; using ‘@SSwarper’).^167,168^ Next, Freesurfer’s ‘recon-all’ was run with default parameters (version 7.0).^174,1758^ This allowed us to segment anatomical images into ventricle and white matter regions of interest to be used to make nuisance regressors.

Preprocessing of the functional data was done using ‘afni_proc.py’^9^ and time series are available as derivatives on OpenNeuro^10^. Blocks were included for despiking (‘despike’), slice-timing correction (‘tshift’), volume registration to the run with the least number of outliers and warping to MNI space (in one step, ‘align’, ‘tlrc’, and ‘volreg’), masking (‘mask’), six mm blurring (‘blur’), scaling to a mean of 100 (‘scale’), and regression (‘regress’). The latter used ‘3dDeconvolve’ to remove demeaned motion and motion derivatives, a polynomial degree of two (‘polort 2’), bandpass filtering (0.01 to 1), ventricular activity (first three principle components), and white matter activity from the time series on a per run basis, in one step. The resulting time series were then edited to account for minor temporal delays caused by movie pauses (on the order of milliseconds). Finally, hand-labelled independent component-based noise regressors were removed from these time series (for details, see).^176^

### Averaging

#### Regression

We used the AFNI program ‘3dDeconvolve’ to conduct a single duration and amplitude-modulated multiple linear regression model on the preprocessed time series to determine the distribution of brain activity for words and their associated sensorimotor properties. This approach allows us to find a ‘main effect’ for words as well as determine how the sensorimotor properties of those words modulate the amplitude of their response. To do this, movies were first annotated for word onset and duration using a combination of machine learning based speech-to-text translation, dynamic time warping to the subtitles, and manual correction (see).^176^ Next, we found the intersection of each word annotation (M = 10,323.80, SD = 2,909.40 words) with the 39,707 words in the Lancaster database of sensorimotor norms.^73^ The norms themselves were created by having participants rate words on 11 sensorimotor experiential dimensions, i.e., auditory, foot/leg, gustatory, hand/arm, haptic, head, interoceptive, mouth, olfactory, torso, and visual.^73^ On average, 95.8% of the words in the movies had a corresponding entry in the database. These were included with the words as 11 sensorimotor word modulators.

We also include three additional ‘nuisance’ modulators to control for the confounding effects of auditory and visual stimulation and word frequency (as used in prior studies).^105,177–179^ Specifically, to account for auditory and visual information that might be correlated with sensorimotor norms in the movie, two ‘low-level’ values were created for each of the words having norms. Sound energy was measured as the root mean square acoustic energy of the movie audio track and was calculated every 100 ms using the Python library ‘librosa’.^180^ Contrast luminance, which measures the standard deviation in luma (brightness) values of the pixels in an image, was computed at every frame in the movie using the Python library ‘OpenCV’.^181^

Both the sound energy and contrast luminance values were averaged over the duration of the words. For instance, if a word had a duration of 200 ms, two values of sound energy (each at 100 ms) would be averaged and 5000 values of contrast luminance would be averaged together (each at 0.04 ms). If word duration was smaller than the sampling rate of either control regressor, the value of one sampling step was assigned to the word. Thus, for instance, if a word had a duration of 90 ms, the sound energy value assigned to the word would be from one 100 ms value, and the contrast luminance would be the average of 2250 values. Across words, the mean sound energy was 8×10^−3^ (SD = 4.5×10^−3^) and the mean contrast luminance was 53.7 (SD = 17.6). Finally, to account for word frequency possibly correlating with specific sensorimotor norms, we created a third ‘nuisance’ modulator value. Because our participants all lived in the United Kingdom, we used the log-transform of word frequency available in the ‘Subtlex-UK’ database.^182^ Only one word did not have an associated word frequency and the mean word frequency across words was 6.2 (SD = 1.2).

In the regression, a block function was used to model the hemodynamic response function whose shape varied by duration and 14 amplitude modulators. These were the 11 sensorimotor norms, contrast luminance, sound energy, and word frequency. Words without sensorimotor norms were modelled with only a duration-modulated block function as a separate regressor (M = 4.2%, SD = 0.8 of words in movies). Finally, we included a duration-modulated regressor for movie times when no words were present (M = 1707, SD = 179 seconds). Regression for words with sensorimotor norms had the following form:

> *word onset * auditory, foot/leg, … visual, sound energy, contrast luminance, word frequency : duration*

#### LME

The resulting words and 11 sensorimotor amplitude modulated beta coefficient maps from the multiple linear regression were input into a linear mixed effects model using ‘3dLME’.^183^ Age, gender, and movie watched for each participant were included as covariates with age centred around the mean. We set participant as a random effect, whereby the intercept of the slope was allowed to vary by a small random amount compared to the group average for each participant. We computed the ‘baseline’ contrasts for words, all 11 sensorimotor norms, and pairwise contrasts between norms.

The results of these contrasts were corrected for multiple comparisons using a multi-thresholding approach (modelled after).^184^ First, we estimated the smoothness and autocorrelation function of neighbouring voxels using the ‘3dFWHMx’ command in AFNI.^185^ Then we ran ‘3dClustSim’ over six uncorrected individual voxel P values (0.05, 0.02, 0.01, 0.005, 0.002, and 0.001) to achieve an alpha (α) threshold of 0.01. Using the significant cluster sizes whereby faces or edges need to touch and voxels are contiguous if they are either positive or negative at each p-threshold, we merged the thresholded maps at each p-threshold to obtain significant voxels (α = 0.01). We discuss and display all results using a minimum cluster size of 20 voxels (540 µL).

We conducted a second duration and amplitude modulated regression and group linear mixed-effects model. This constituted an additional analysis to try and understand the lack of inferior frontal gyrus activity for words in the analysis described in this and the prior section. We reasoned that word frequency likely accounts for the lack of activity given it has been directly demonstrated to increase activity in these regions with decreasing frequency.^74–76^ The analyses were the same as described but did not include the sensorimotor modulators. To have something to compare to, we contrasted word frequency directly with sound energy. This also allowed us to determine whether sound energy accounted for activity in ‘lower-level’ auditory-related brain regions, providing reassurance that it was serving its role as a nuisance regressor.

#### Hubs

We next sought to determine if the voxels corresponding to the main effect of words in the 3dLME model correspond to hubs. We did this using measures of centrality, how important a node is for the integrity and information flow within a network. Centrality can be determined using various metrics that provide different information on the role of the node of interest in the network. We measured degree, eigenvector, closeness, and betweenness centrality to ensure that we are robustly characterising centrality across metrics.^48,186–188^

Degree centrality is the sum of inward and outward connections from a node. Eigenvector centrality is a measure of influence on a network, meaning that a high-connectivity node linked to nodes of high connectivity will have higher eigenvector centrality (i.e., be more influential) than a high-connectivity node linked to low-connectivity nodes. Betweenness centrality measures the shortest paths that pass through a given node and closeness centrality measures the inverse of the distance of shortest paths passing through the node. Although these centrality metrics provide different details on the importance of a node, they are correlated to one another.^188,189^

We calculated these four measures of centrality on a voxel-wise basis and in a dynamic manner. To do this, we constructed time-varying connectivity matrices using a sliding-window approach. First, the time series was resampled from three to a four mm^3^ resolution (using ‘3dresample’) due to the lengthy time calculated for analyses at the native three mm resolution, considering the available computational resources. Resampled time series were then divided into windows of one-minute length, sliding every five seconds to allow for a 55-second overlap between one window and the subsequent window. Next, a pairwise Pearson’s product-moment correlation coefficient was computed on a voxel-wise basis at each window using the AFNI program ‘3dDegreeCentrality’.^185^ Finally, the resulting correlation matrices were proportionally thresholded to obtain a 5% sparsity in each time window. These values were considered a connection between two voxels and used to build connectivity matrices which were used to calculate the centrality values described.

Despite some differences, these four centrality metrics had a significant Spearman’s ranking correlation with one another at the group level (i.e., across time windows and participants; M_ρ_ = 0.94, SD_ρ_ = 0.02; p ≤ 0.001). Thus, to simplify analyses and conform to our hubs/periphery hypotheses, we clustered the four centrality values using Ward’s minimum variance,^83^ which consistently divided voxels in each time window into two groups (M = 2.00, SD = 0.02) across all participants, one high-centrality (labelled 1) and one low-centrality cluster (labelled 2). In rare cases, the clustering method detected greater than two clusters of centrality, but since the vast majority of windows divided centrality values into two groups, we recomputed the few outlier time windows by forcing them to split the data into two clusters to investigate spatial variations in connectivity over time.

Next, we divided the resulting windowed centrality time series into a high and a low centrality time series for each participant and averaged across all windows. The resulting maps were used to conduct a linear mixed effects model with ‘3dLME’ (described above).^183^ The fixed effects were centrality (high and low), age, gender, and movie with participant as a random effect. A correction for multiple comparisons using a cluster-size correction with multiple voxel-wise thresholds was applied (again, as described above, using α = 0.01).

We then submitted the full windowed high and low centrality time series from all participants to a group spatial independent component analysis (ICA) using the default parameters of ‘MELODIC’ (version 3.15), reducing to a 100-dimensional subspace.^190^ These results were then used in a dual regression to estimate a version of the 100 resulting components for each participant so group statistics could be calculated. Specifically, this approach first regresses the group spatial maps into each participant’s time series to give a set of timecourses. These timecourses are then regressed onto the time series to get 100 participant-specific spatial maps for both high and low centrality. These were then directly contrasted using a t-test and thresholded using a voxel-wise z-value equivalent to p ≤ 0.0001 (i.e., 0.01 divided by the number of components) and a minimum cluster threshold of 20 voxels (540 µL) to protect against multiple comparison issues. Finally, we computed the spatial correlation of each of these contrasts with words from the above 3dLME analysis to locate a hub or hubs corresponding to words.

We also averaged each time window across participants for *500 Days of Summer* and *Citizenfour* separately to conduct two additional MELODIC ICAs, again with 100 dimensions, with resulting components thresholded using a mixture modelling approach.^190^ We did not do this on the other eight movies because they had only six participants each. We also conducted a similar ICA analysis on the centrality maps directly thresholded at 90% to define high centrality regions across all 86 participants. This was done to provide evidence that results were not dependent on the more categorical high/low centrality clustering method used for all of the primary analyses. All other analyses are as described in the Results section. For display purposes, we also conducted affinity propagation clustering as described in the Supplementary Fig. S2 and S3 captions.^191,192^

### Connections

Next, we evaluated whether linguistic representations are preferentially encoded in dynamic connectivity between ‘language hubs’ and peripheral regions or in the regions themselves. To do so, we developed encoding models that mapped contextual language embeddings onto voxel-level activity and inter-regional connections, based largely on prior work with regional activation.^84,85^

This analysis was first conducted on the *500 Days of Summer* movie-fMRI data (N = 20). It was applied to two masks derived from the LME results described in the ‘Averaging’ section: the ‘language hub’ mask, defined by binarising significant voxels from the baseline contrasts for words (see Fig. 5, black outline); and the periphery mask, derived by binarising all significant voxels from the 11 sensorimotor norms and pairwise contrasts between them (see Fig. 5, hot colours).

To align language model embeddings with fMRI data, we tokenized the full stimulus transcript using the GPT-2 tokenizer implemented via the *HuggingFace* Transformers library (v4.40). We used the pretrained GPT-2 small model, which consists of 12 transformer layers and approximately 117 million parameters. Hidden-state representations were extracted from layers 8 and 12, which have been suggested to encode intermediate and late-stage semantic and syntactic information.^85,114^ For each token in the transcript, we extracted its hidden-state vector from the relevant layer, yielding one 768-dimensional vector per token. These vectors were then temporally aligned to the fMRI data (following^85^). Specifically, the full sequence of token embeddings was divided into T contiguous and equally sized bins, where T is the number of fMRI volumes (TRs) collected during the scan. For each bin, we summed the hidden-state vectors of all tokens it contained, producing one embedding vector per TR. If a bin contained no tokens (e.g., due to pauses), we inserted a zero vector, preserving the one-to-one correspondence between embedding vectors and fMRI volumes. This yielded a final matrix of shape (T, 768), representing the time-resolved linguistic embedding of the movie.

To account for the temporal dynamics of the hemodynamic response, we expanded this TR-aligned embedding sequence into a temporally lagged design matrix by concatenating six shifted versions of the embedding sequence (lags 0 to 5). This produced a matrix of shape (T, 768 × 6), where each row contained information from the current and five preceding timepoints. The temporal design was constructed using a custom *PyTorch*-based implementation that efficiently shifts and concatenates the embeddings into a single feature matrix. This representation was used to model both voxelwise activation and dynamic connectivity patterns.

To model the dynamic coupling between brain regions and linguistic input, we implemented a 10-TR sliding window approach applied to the fMRI data. At each TR, a centered window of 10 timepoints (padded via reflection at the boundaries) was extracted from voxel time series within the hub and periphery masks. Within each window, Pearson correlations were computed between the time series of each hub voxel and all voxels in the periphery mask. These correlations were weighted using a Hanning kernel to emphasize central timepoints and reduce edge artifacts. The resulting pairwise correlations were averaged across periphery voxels to yield a single dynamic connectivity value per hub voxel per TR, forming a time-resolved matrix of functional coupling. In parallel, we extracted voxelwise activation time series by reshaping the 4D fMRI volumes into matrices of shape (T, V), where V is the number of voxels in either the hub or periphery mask. These activation traces were modeled using the same temporally lagged GPT-2 design matrix, enabling the prediction of both local activation and regional connectivity directly from the stimulus-derived embeddings.

To estimate encoding weights, we used a combination of custom and standard tools. A *PyTorch*-based implementation constructed the temporally lagged design matrix, and *RidgeCV* from *scikit-learn* was used to fit a ridge-regularized linear model while automatically selecting the optimal regularization strength. Model selection followed a nested cross-validation procedure: a five-fold outer loop with shuffled timepoints to assess generalization, and a three-fold inner loop to select the best α value from {0.01, 0.1, 1, 10}. For each fold, regression weights were estimated using the closed-form solution:

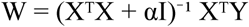

where X is the lagged embedding matrix and Y is the target signal—either voxelwise activity or hub–periphery connectivity time series. Voxelwise models predicted the activity of each voxel independently; connectivity models predicted the dynamic coupling time series for each hub voxel. Predictions were evaluated using Pearson correlation between predicted and observed time series, computed per voxel or connection and per fold.

While voxelwise models captured univariate activation patterns in localized regions, connectivity models captured coordinated fluctuations between regions, offering a complementary view of functional integration. The same voxel masks were used across all windows, with spatial definitions held constant; only the temporal context for computing connectivity was dynamic. Model performance was assessed using five-fold temporal cross-validation. Prediction accuracy on held-out timepoints was quantified using Pearson correlation, Fisher z-transformed, and averaged across folds. For each participant, we computed the mean positive Fisher z value across voxels or connectivity targets for each condition (hub activation, periphery activation, hub–periphery connectivity) and for each model layer.

For statistical testing, we performed paired two-tailed t-tests comparing mean Fisher z scores across the three encoding conditions, testing the hypothesis that hub–periphery connectivity encodes linguistic information more robustly than regional activation alone. All statistical tests were performed using *SciPy* (v1.11). All modeling, evaluation, and visualization procedures were implemented in Python 3.10 using custom scripts and standard scientific computing libraries.

#### Audiobook-fMRI

We replicated central movie-fMRI analyses using an fMRI dataset of participants listening to an audiobook to test whether movie-related findings generalize to audio-only stimulation.^70^ Specifically, we analyzed data from 49 English-speaking participants (30 female, mean age = 21.3 years, SD = 3.6) who listened to *Le Petit Prince* (94 minutes) during 3T multi-echo fMRI scanning. Preprocessing was initially performed by the dataset authors using AFNI. We further applied spatial smoothing, scaling, and drift correction where needed (also using AFNI). The first 100 seconds of data were excluded due to accompanying static images. All analyses were approximately the same as in the movie-fMRI, except that the multiple linear regressions excluded a luminance modulator, the group analysis excluded the ‘movie’ covariate, and the encoding model analysis used 20 second windows. Full details are available in the ‘Audiobook-fMRI’ section of the Supplementary Materials.

#### sEEG

Experimental data were recorded from 10 adult patients without known language impairment undergoing intracranial monitoring for epilepsy (ages 22-45; 6 males and 4 females). All procedures were approved by the institutional review board (IRB) at Baylor College of Medicine (BCM). Reported data were acquired by stereo stereoelectroencephalography (sEEG) depth probes. The sEEG probes had a 1.28 mm diameter, with nine recording contacts, with a 5.0 mm centre-to-centre distance between contacts (AdTech Medical Instrument Corporation).

Local field potential (LFP) signals were recorded by a 256 channel BlackRock Cerebus system (BlackRock Microsystems) at 2 kHz sampling rate, with a fourth order Butterworth bandpass filter of 0.3–500 Hz. SEEG recordings were referenced to a reference electrode placed in the subgaleal space. All signals were processed in MATLAB (R2023b, MathWorks, MA, USA). Raw sEEG signals were first inspected for line noise, recording artefacts (e.g., contamination by muscle activity, presence of large deflections, additional noise components) and interictal epileptic spikes. Electrodes with artefacts and epileptic spikes were excluded from further analysis. Next, the signal from each ‘clean’ electrode was notch filtered (60 Hz and harmonics) and each electrode was re-referenced to a common average of all clean electrodes. Finally, re-referenced signals were spectrally decomposed using a family of Morlet wavelets (seven cycles), with center frequencies ranging logarithmically from 1 to 200 Hz in 100 steps. Frequency band power was obtained by averaging the magnitude of the Morlet wavelet decomposition result between 70–150 Hz for broadband gamma.

For each patient, we determined electrode locations by employing the software pipeline intracranial Electrode Visualization, iELVis.^193^ In short, the postoperative CT image was registered to the preoperative T1 anatomical MRI image using FSL.^194^ The location of each electrode was then identified in the CT-MRI overlay using BioImage Suite.^195^ The electrode anatomical locations were classified based on their proximity to the cortical surface model, reconstructed by the T1 image using Freesurfer (version 6.0).^196^ The anatomical assignment of each electrode was manually verified by an expert in neuroanatomy (Bachelor of Science). We further labelled each electrode by mapping an atlas^197^ onto each individual cortical surface and finding the most likely cortical parcellation estimate using a 4 mm radius around each electrode. To visualize electrodes of interest on an example patient brain, we represented them as spheres using the Multi-Modality Visualization Tool, MMVT^198^ (Fig. 7a). For each patient, electrodes were classified as ‘language’ or ‘non-language’ by comparing their MNI coordinates to the ‘language’ region coordinates in the large-scale meta-analysis software Neurosynth (also used in Fig. 2).^165^ Electrodes in white matter were not included in the analysis. Across ten patients, we had 544 total electrodes in ‘language regions’, ranging from 23–110 for each patient, and we had 1,001 total electrodes in ‘non-language regions’ (including the amygdala, thalamus, putamen, insula, hippocampus, right hemisphere temporal cortex and prefrontal regions—anterior cingulate, orbitofrontal cortex, and medial PFC), ranging from 61-143 for each patient.

### Stimuli

Patients listened to six 5- to 13-min stories taken from The Moth Radio Hour, totaling 45 minutes of listening time^199^. The six stories were, ‘Life Flight’, ‘The Tiniest Bouquet’, ‘The One Club’, ‘Wild Women and Dancing Queens’, ‘My Father’s Hands’, and ‘Juggling and Jesus’. In each story, a single speaker tells an autobiographical story in front of a live audience. The six selected stories were chosen to be both interesting and linguistically rich. Stories were played continuously through the built-in audio speakers of the patient’s hospital TV room. The audio signal is synchronized to the neural recording system via analog input going directly from the computer playing the audio into the Neural Signal Processor at 30kHz. After the experiments, the audio .wav file was automatically transcribed using Python and Assembly AI^11^, a state-of-the-art AI model to transcribe and understand speech. The transcribed words and corresponding timestamp output from Assembly AI was converted to a TextGrid and then loaded into Praat, a software for speech analysis. The original .wav file was also loaded into Praat and the spectrograms and timestamps were manually checked and corrected to ensure the word onset and offset times are accurate. The TextGrid output of corrected words and timestamps from Praat was converted to a xls and loaded into Matlab and Python for further analysis. This stimulus contained 7,346 words that were used in the analysis.

To extract part-of-speech (POS) for each word in the dataset, we utilized an automated pipeline through Stanford CoreNLP, a natural language processing toolkit.^200^ We initialized a CoreNLPParser with the ‘pos’ tagtype, which specializes in part-of-speech tagging. The transcript was first segmented into sentences based on punctuation. Each sentence was then tokenized and passed through the

CoreNLPParser’s tagging function. This process leveraged CoreNLP’s advanced linguistic models to analyze the context and structure of each sentence, assigning appropriate POS tags to individual words. The 22 POS types were: Nouns, pronouns, verbs, adjectives, adverbs, determiners, articles, prepositions, conjunctions, interjections, numerals, particles, modal auxiliaries, possessive endings, ‘to’ as a preposition or infinitive marker, wh-words, existential ‘there’, foreign words, list item markers, proper nouns, predeterminers, and possessive wh-pronouns. This dataset contained 1,234 nouns, 1,466 verbs, and 4,646 other POS words that were used in analysis.

### Decoding

For each word, gamma power from each electrode was summed across word duration and divided by word duration. We used a SVM decoder^201^ with a linear kernel to determine whether the gamma power during words carries information about syntax – specifically nouns and verbs. We computed gamma power on each electrode for all nouns and verbs and classified binary labels specifying the event (for example, verbs were class one, nouns were class two) from these neural data. Electrodes in ‘language regions’ (as defined by meta-analysis, see above) were pooled across all patients to create a word by electrode matrix known as the pseudopopulation (Note that, results shown in Fig. 7c are consistent when computed for each patient separately). We also compiled the pseudopopulation for ‘non-language’ electrodes and modeled them separately. The number of observations of nouns and verbs were always balanced. Since there were more electrodes in ‘non-language regions’ (1,001 electrodes), we randomly subsampled the number of these electrodes to match the number of ‘language’ electrodes (554 electrodes) on each iteration of model training. Random selections of class observations were repeated for 100 iterations, giving us the average classification accuracy over 1,000 test splits of the data for each session. To train and test the model, we used a tenfold cross-validation. In brief, the data were split into ten subsets, and in each iteration the training consisted of a different 90% subset of the data; the testing was done with the remaining 10% of the data. We used the default hyperparameters as defined in *fitcsvm*, MATLAB 2023b, with z-scored normalization of gamma power across electrodes. Decoder performance was calculated as the percentage of correctly classified test trials. We compared model performance for predicting train and test data to check for overfitting. In each iteration, we trained a separate decoder with randomly shuffled class labels. The performance of the shuffled decoder was used as a null hypothesis for the statistical test of decoder performance.

POS encoding of each electrode during words was quantified using an ROC (receiver operating characteristic) analysis, a standard approach that has previously been used to characterize neural responses during behavior.^202,203^ Here, the gamma power on each electrode is the classifier that is evaluated on its ability to distinguish nouns from all other words (except verbs), and vice versa. Upon application of a binary threshold to gamma power and comparison with a binary event vector denoting word label (1 for noun, 0 for other POS), POS detection based on neural activity can be measured using the true positive rate (TPR) and the false positive rate (FPR). Plotting the TPR against the FPR over a range of binary thresholds, spanning the minimum and maximum values of the neural signal, yields an ROC curve that describes how well the neural signal detects POS at each threshold. With the *perfcurve* function in Matlab 2023b, we used the area under the ROC curve (AUC) as a metric for how strongly electrodes are modulated by each word POS (Fig. 7d). For each electrode and event category (nouns and all other, verbs and all other), the observed AUC was compared to a null distribution of 1,000 AUC values generated from constructing ROC curves over randomly permuted neural signals (that is, gamma power permuted using a random time shift across words, thus destroying the relationship between neural data and POS labels). The P value is calculated as the proportion of permuted AUC values that are greater than or equal to the observed AUC and an electrode was considered significantly responsive if P < 0.05. This analysis was performed twice, once for nouns and other words, and again for verbs and other words, for electrodes in ‘language regions’ and electrodes in ‘non-language regions’. The number of words in the Noun and Verb class were balanced with the number of words in ‘Other’ class by subsampling words from the ‘Other’ class over 100 iterations of the analysis. For each electrode, AUC was computed, then averaged and compared across iterations.

### Hubness

For ‘language’ and ‘non-language regions’ separately, the AUC values from respective electrodes for both nouns and verbs were combined into one vector and used in principal component analysis (pca, Matlab 2023b). The output variable, ‘latent’, values reflect the eigenvalues or amount of variance that each principal component explains in the dataset. We divided latent 2 by latent 1 to compute the ratio (dashed line) shown in Fig. 7e, from noun-verb ROC values for ‘language’ and ‘non-language regions’. To compare ratios between ‘language’ and ‘non-language regions’, we generated ratio distributions by resampling the original vector of AUC noun and verb values with replacement 1,000 times. The P value reported in Fig. 7e is computed from the Wilcoxon rank-sum test between these two bootstrapped distributions.

## Acknowledgements

This work was partially supported by a doctoral training grant given to SA from the UK Research and Innovation (UKRI) Biotechnology and Biological Sciences Research Council (BBSRC) as part of the London Interdisciplinary Biosciences Consortium (LIDo). We would like to thank the Birkbeck-UCL Centre for Neuroimaging (BUCNI) for supporting the NNDb. We thank the Texas Advanced Computing Center (TACC), especially Christopher Simmons, Georgia Stuart, and Yile Wang for helping with computation resources. BH is supported by the McNair Foundation. We thank The LAB Lab more generally for continued discussion. JIS is supported by the Wellcome Leap and would like to thank the patience of a special Banana.

## Author Contributions

JIS and SA conceived of the study. JIS did the fMRI preprocessing. SA developed and did NNDb analyses herein with *500 Days of Summer* and wrote the first draft of the manuscript. SA, BW, and JIS extended these initial analyses to the other nine NNDb movies and 66 participants. JIS did all meta-analyses work. AG and GC conducted the replication analyses with the audiobook-fMRI dataset. JIS and VK conceived of and VK carried out the Connections encoding model analysis. MF and BH conducted the LFP work. JIS wrote the final manuscript and made all of the figures, except the LFP figure, made by MF. SLS was involved in conceptualisation and writing.

## Open Science Statement

This study was not preregistered. A version of this work is available in SA’s Doctoral thesis, ‘A Model of the Network Architecture of the Brain that Supports Natural Language Processing’. A version of this paper is available as a preprint on bioRxiv (https://www.biorxiv.org/content/10.1101/2023.09.01.555886). The preprocessed data used in analyses is publicly available on OpenNeuro (https://openneuro.org/datasets/ds002837/versions/2.0.0). The code used to generate results will be made available on GitHub (https://github.com/lab-lab/).

## Footnotes

1 ‘Language regions’ and ‘language network’ result in more than 12,000 and 26,000 links on Google Scholar, respectively (ascertained July, 2025).

2 For example, to Bonferroni correct a single ‘mass univariate’ statistical image with two mm^3^ voxels, the P value should be 0.05/250,000 or p ≤ 0.0000002. That said, corrections are often not this stringent because allowances are made for the fact that voxels are far from independent.

3 Thus, the word ‘averaging’ as used throughout is a convenient simplification for this particular set of analyses steps and other similar neuroimaging analyses. Arguably, all neuroimaging studies involve measures of central tendency or averaging in this sense.

4 Note that all calculations done in this paragraph are similar when done with the language meta-meta-analysis instead of the main effect of words.

5 Note that results need not have turned out this way, e.g., word regions could have been consistently low centrality regions.

6 Calculated using https://neurovault.org/images/57681/

7 https://afni.nimh.nih.gov/

8 http://www.freesurfer.net/

9 https://afni.nimh.nih.gov/pub/dist/doc/program_help/afni_proc.py.html

10 https://openneuro.org/datasets/ds002837/versions/2.0.0

11 https://www.assemblyai.com/

