## Supplementary Materials for "*The entire brain, more or less, is at work*: ‘Language regions’ are artefacts of averaging"

#### Illustration

To illustrate the theoretical results of averaging in neuroimaging studies of language (Fig. 1), we defined three sets of regions: a set of putative ‘language regions’ and two complementary sets of putative ‘non-language regions’ related to sensory/motor and cognitive/emotional processing. The ‘language regions’ were defined using a set of left hemisphere parcels created from a probabilistic overlap map of 220 participants’ language localizer activation maps, subsequently mirrored to the right hemisphere, and currently in use by ‘EvLab’ (see[1]). From this map, we randomly selected nine out of 12 parcels as our working set of ‘language regions’ (Fig. 1, red).

The ‘non-language regions’ were derived from the 200-region, 7-network Schaefer cortical parcellation atlas[1] (see also[2]), resampled to match the spatial resolution of the language localizer map. We manually classified these parcels into two broad functional groups. The sensory/motor group consisted primarily of parcels associated with visual, somatomotor, and select salience-related regions, while the cognitive/emotional group included dorsal attention, limbic, frontoparietal control, and default mode network parcels. To ensure mutually exclusive sets, any parcels overlapping with the defined ‘language regions’ were excluded from the Schaefer atlas. Our classification was intentionally coarse, designed to ensure whole-brain coverage and balanced group sizes (62 parcels for sensory/motor; 70 for cognitive/emotional), rather than strict anatomical or functional specificity. The classification was ultimately arbitrary and used to construct contrasting ‘non-language’ processing

[1] https://www.evlab.mit.edu/resources-all/download-parcels
[2] https://github.com/ThomasYeoLab/CBIG/tree/v0.14.3-Update_Yeo2011_Schaefer2018_labelname/stable_projects/brain_parcellation/Schaefer2018_LocalGlobal/Parcellations

systems for illustrative purposes only; the precise composition of these groupings should not affect the general outcome of the simulation.

To simulate co-activation patterns, we generated 10,000 iterations in which 15 regions were randomly sampled per iteration: nine from the ‘language regions’ (Fig. 1, red), three from the sensory/motor group (Fig. 1, green), and three from the cognitive/emotional group (Fig. 1, blue). Each set of regions was binarized and projected from volume space onto the *fsaverage* cortical surface (fetched with *nilearn.datasets.fetch_surf_fsaverage*) using volumetric-to-surface interpolation via *nilearn.surface.vol_to_surf*. The frequency of voxel selection across iterations was aggregated, producing a probabilistic spatial map of region involvement. These data were visualized in aggregated form, thresholded at the 90th percentile.

To complement this regional simulation, we constructed synthetic connectivity-like patterns using adjacency matrices. Each of the 10,000 iterations began with the same set of 144 nodes: 12 ‘language regions’, 62 sensory/motor, and 70 cognitive/emotional regions. Within each iteration, we simulated sparse inter-regional connections as follows: one randomly selected pair of ‘language regions’ was assigned a bidirectional connection; one ‘language region’ was connected to three randomly selected sensory/motor regions; and one ‘language region’ was connected to three randomly selected cognitive/emotional regions. This produced a symmetric adjacency matrix per iteration, which we accumulated and normalized to generate an average connectivity matrix. We then applied a 90th-percentile threshold (of nonzero values) to retain only the strongest connections, which were further normalized and scaled using a power-law exponent to enhance visual contrast.

Connection matrices were visualized using the *pycirclize* package, which generates chord diagrams representing node-to-node relationships in circular space. Each node was labeled and color-coded by group in Fig. 1: red for ‘language regions’, green for sensory/motor, and blue for cognitive/emotional regions. We generated example connectivity plots from randomly selected iterations and one final plot summarizing average connectivity across all iterations. Edge thickness in the chord diagrams was scaled according to normalized connection strength.

#### Audiobook-fMRI

We replicated key movie-fMRI results with audio-only audiobook-fMRI to evaluate whether they are the product of visual stimulation.

#### *Methods*

We used an fMRI dataset derived from 112 participants who listened to the audiobook *Le Petit Prince* in their native language during multi-echo fMRI acquisition.[2] Analyses were limited to the English-speaking participants (n = 49, 30 females, mean age = 21.3 years, SD = 3.6 years). All participants were right-handed, paid, and gave written informed consent in accordance with the IRB guidelines of Cornell University.

Per the authors of the original study, functional MRI scans were collected on a 3T MRI GE Discovery MR750 using a 32-channel head coil and a multi-echo EPI sequence (TR = 2 seconds, TEs = 2.8, 27.5, 43ms, flip angle of 77°, 33 axial slices, in plane resolution = 3.75mm x 3.75mm, slice thickness = 3.8mm, 2x image acceleration). The English audiobook is 94 minutes long, which was divided into 9 runs of ~10 minutes each. Anatomical scans were collected using a T1W MPRAGE pulse sequence (resolution = one mm).

Initial preprocessing was performed by the dataset creators using the AFNI software package. Anatomical images were deskulled and nonlinearly aligned to the MNI N27 template brain. Functional image preprocessing included slice-timing correction (3dTshift), despiking (3dDespike), volume registration (3dvolreg), and nonlinear alignment to the MNI N27 template brain. Multi-echo ICA was used to remove motion, physiological, and scanner artifact noise, and images were resampled (3dresample) to 2mm cubic voxels.

In addition to these, we performed further preprocessing steps. These included blurring the functional data to six mm (using 3dBlurToFWHM) and scaling to a mean of 100 (3dTstat and 3dcalc). For functional connectivity analyses, we also used 3dDetrend to remove slow drift and baseline trends using a polynomial order of five, following the AFNI rule of thumb where polort = 1 + floor(run duration in minutes / 150). Finally, because the first 100 seconds of the audiobook were presented to participants with static visual images, we removed these time points to ensure our analyses were limited to audio-only stimulation.

We replicated most analyses from the NNDb movie-fMRI dataset on this audio-only dataset. For the duration and amplitude-modulated multiple linear regression (using ‘3dDeconvolve’), we used the word annotations, onsets, offsets, sound intensities, and word frequencies provided by the dataset creators. Duration was calculated by subtracting each word’s onset from its offset. The dataset authors provided an initial list of lemmatized versions of the words, and we further lemmatized the list using the WordNetLemmatizer function in Python’s Natural Language Toolkit (e.g., the first clause “once when i was six years old” became “once when i be six year old” after our additional lemmatization process). This allowed us to find Lancaster sensorimotor norms for 97.6% of the words in the audiobook (15,011/15,376 words with norms). Regression for words with sensorimotor norms followed the form of our movie-fMRI regressors with the exception of a contrast luminance modulator; thus, the form was:

*word onset * auditory, foot/leg, ... visual, sound intensity, word frequency : duration*

Words without sensorimotor norms were modeled with a separate duration-modulated block function including modulators for sound intensity and word frequency. Times when no words were present, which occurred at the end of each run, were included as a duration-modulated regressor without additional modulators. The individual-level results from the regression were analyzed at the group-level in concordance with the linear mixed effects model used on the movie-fMRI data, with the exclusion of the covariate ‘movie watched’ because all participants listened to the same audiobook. The results were multi-thresholded with the same P values as the movie-fMRI results, to achieve $\alpha < 0.01$, and presented with a minimum cluster size of 20. All ‘Hubs’ and ‘Connections’ analyses were performed using the same code as for the movie-fMRI, modified to accommodate the audiobook-fMRI file names and directory structure.

Finally, unlike the movie-fMRI analysis, the encoding model analysis was conducted using all 49 participants from the audiobook dataset. We used the same analysis parameters without adjusting the window length. Because the TR was 2 seconds rather than 1, each sliding window spanned 20 seconds rather than 10. We view this as a strength: the fact that results replicated despite differing temporal resolution suggests they are robust to variation in timeseries parameters.

#### *Results*

##### Averaging/Distributed

As with movie-fMRI, sensorimotor processing was distributed throughout much of the brain during audiobook-fMRI (Fig. S5, yellows and reds). Fig. S5 shows the main effect of the 11 sensorimotor modulators (e.g., foot/leg, Fig. S5, reds) and pairwise contrasts between modulators (e.g., foot/leg vs. hand/arm, Fig. S5, yellows) averaged and projected onto the brain, along with voxels that were not significant in any of these comparisons (Fig. S5, blues). Numerically, the audiobook-fMRI results closely matched the movie-fMRI results. In total, 37.75% (compared to 36.47% for movie-fMRI) of the brain was activated by sensorimotor modulators outside of voxels activated by words (483,016 of 1,279,432 μL), with each modulator activating an average of 4.99% (compared to 4.32% for movie-fMRI) of those voxels (63,858.91 of 1,279,432 μL, SD = 40,656.25). Similarly, when considering direct contrasts between modulators, 67.66% (compared to 66.88% for movie-fMRI) of the brain was activated outside of word voxels (865,688 of 1,279,432 μL), with each contrast activating an average of 6.17% of those voxels (compared to 4.31% for movie-fMRI; 78,901.38 of 1,279,432 μL, SD = 52,433.73). For comparison, words activated about 10.86% (compared to 4.57% for movie-fMRI) of the whole brain in total (155,864 of 1,435,296 μL). Finally, neither the spatial pattern of activity for the unthresholded sensorimotor (mean r = 0.05, SD = 0.46; compared to r = -0.02 for movie-fMRI) nor the direct contrasts maps (mean r = 0.02, SD = 0.44; compared to r = -0.05 for movie-fMRI) were correlated with the unthresholded word map on average.

**Hubs**

As with movie-fMRI, results with the LME show that high centrality voxels are significantly more active than low centrality voxels in much of the superior and middle temporal plane, (Fig. S6, colours) and that the main effect of words overlap with these regions (Fig. S6, black outline). Indeed, 78.74% (versus 78.26% for the movie-fMRI) of the word voxels were high > low centrality (thresholded at $\alpha$ = 0.01 with a minimum individual voxel P value $\leq$ 0.001). Even when thresholding results further to include only the top 90% of the high centrality voxels, there was still a 21.51% overlap (compared to 39.00% for movie-fMRI).

Next, we determined whether regions delineated by words formed a coherent set of hubs, independent of other sets in the aggregate across time. To do so, we again performed group spatial independent component analysis (ICA) with 100 dimensions across participants' dynamic high and low centrality time series. We then used dual regression to quantify the differences in high and low centrality, involving a direct contrast and thresholding using a correction for the 100 comparisons conducted. Finally, we computed the spatial correlation of each of the resulting contrasts with words, requiring a moderate or larger threshold (i.e., r $\geq$ 0.30). Results reveal one contrast correlated with words (IC 11, r = 0.47; two additional ICs survive at r $\geq$ 0.10, i.e., IC 6, r = 0.10 and IC 9, r = 0.14; similar to the movie-fMRI results). This IC was significantly more associated with high than low centrality (Fig. S6, black outline; compare to Fig. 6, shaded white outline). IC 11 was associated with the transverse temporal gyrus and the superior temporal gyrus, bilaterally. Including ICs 6 and 9, largely fills out the white main effect of words outline, particularly in the posterior aspects.

Thus, as with movie-fMRI, both the linear mixed effects and dual regression analyses suggest that voxels associated with words form a coherent set of connectivity hubs. Results also suggest that these only appear in the aggregate over time and do not exist as a whole on a moment-to-moment basis. That is, hubs are not fixed but are dynamic, e.g., moving around the superior and middle temporal lobes to coordinate distributed and varying peripheral regions. If this is the case, individual time windows should not be correlated with words as highly as in the ICA analysis. To examine this, we averaged across participants for each window for our audiobook. We thresholded the voxels in the resulting time series at 90% (i.e., $\geq$ 1.8) of the mean maximum value (i.e., 2) to isolate high centrality voxels. We then spatially correlated each thresholded time window with words. The mean correlation of the audiobook was $r = -0.02$ ($SD = 0.02$; compared to $r = 0.03$ for ‘500 Days of Summer’ and $r = 0.02$ for for ‘Citizenfour’) with no time window having a correlation $\geq 0.10$ (compared to $\leq 1$ time window for $r \geq 0.30$ and $\leq 11\%$ of time windows for $r \geq 0.10$ for the two movies). Performing ICA on the high centrality audiobook timeseries produced similar results as those in the prior paragraph, with one network correlated with words $r \geq 0.30$ (IC 24, $r = 0.41$; compared to 2 networks for each of the two movies).

Finally, it might be the case that highly central word hubs are not connected to the sensorimotor periphery, e.g., if the periphery is ‘extraneous’ and processing is proceeding there independently. To address this, we calculated the specific connectivity profiles of windows on an individual participant basis where the mean cluster assignment of the voxels from the word mask was $\geq$ 90% (i.e., $\geq$ 1.8, returning to the more categorical high/low windows) to the periphery defined, again, as the sensorimotor maps (i.e., the connectivity between the regions within the black outline and those in colour in Figs. 2-5). This analysis revealed that when the word mask was a hub, it shared an average of 38.24% ($SD = 4.44$; Range = 0.00 to 49.74%; Compared to Mean = 43.99% and $SD = 1.33$ for movie-fMRI) of connections with peripheral voxels across all participants. This helps explain the low spatial correlation between the thresholded windows and the word map and the relatively low percentage of word masks acting as hubs at any given time window above. That is, when word voxels are hubs at any given moment, they are connected to a periphery, driving down correlations with the word map that has been aggregated over time, thus excluding the less central and more variable periphery.

**Connections**

Group-level paired two-tailed t-tests on participants’ mean positive Fisher-z values replicated results from the movie-fMRI analysis. Overall, voxel-wise prediction accuracies varied significantly across conditions. Specifically, connectivity patterns between core hubs and peripheral regions outperformed both hub and peripheral activations alone, with peripheral activations consistently yielding the lowest prediction accuracy. At layer 8, connectivity (mean Fisher-z = 0.08) was approximately 4 times more accurate than hub activations (mean = 0.02; $t = 10.87$, $p < 0.001$) and 8 times more accurate than peripheral activations (mean = 0.01; $t = 10.78$, $p = 0.85$). Similarly, at layer 12, connectivity (mean Fisher-z = 0.11) was about 5.5 times more accurate than both hub (mean = 0.02; $t = 21.58$, $p < 0.001$) and peripheral (mean = 0.02; $t = 19.9$, $p < 0.001$) activations. These findings reinforce the idea that linguistic information is best captured by connectivity between language hubs and the broader brain periphery. For a summary of results, see Fig. S4.

### Supplementary Figures

#### *Figure Captions*

**Fig. S1. Contrast of word sound energy and frequency nuisance modulators.**

For comparison, the white outline corresponds to the language meta-meta-analysis (see Fig. 1) and the black outline corresponds to the main effect of words (see Fig. 4). All results were cluster-size corrected for multiple comparisons at $\alpha = 0.01$ and displayed with a minimal cluster size of 20 voxels (540 μL).

**Fig. S2. Independent components from dual regression of high and low centrality correlated with words.**

We conducted independent components analysis using a dimensionality of 100 and performed a direct contrast between high and low centrality with a t-test following dual regression. We clustered the resulting unthresholded spatial independent components (ICs) using affinity propagation clustering. We present only the part of the dendrogram with the ICs that correlated with the main effect of words at $r \geq 0.30$ (right, under the red lines/text; ICs with $r \geq 0.10$ and $r \leq 0.30$ are indicated under the blue lines/text). The full dendrogram is on the left and the site where these ICs are located is indicated with red lines on that dendrogram. The axial and sagittal brain images are from the centre of mass of the largest cluster in each IC. These images are thresholded at $\alpha = .01$; $p \leq 0.0001$ with a minimum cluster threshold of 20 voxels (540 μL).

**Fig. S3. Independent components analysis on high centrality regions defined as >90% of the average of the z-transformation of four centrality values correlated with words.**

We conducted independent components analysis using a dimensionality of 100. We clustered the resulting unthresholded spatial independent components (ICs) using affinity propagation clustering. We present only the part of the dendrogram with the ICs that correlated with the main effect of words at $r \geq 0.30$ (right, under the red lines/text; ICs with $r \geq 0.10$ and $r \leq 0.30$ are indicated under the blue lines/text). The full dendrogram is on the left and the site where these ICs are located is indicated with red lines on that dendrogram. The axial and sagittal brain images are from the centre of mass of the largest cluster in each IC. These images are thresholded using a mixture modelling approach with a minimum cluster threshold of 20 voxels (540 μL).

**Fig. S4. Encoding model accuracies.**

Mean voxel-wise encoding accuracies (Fisher-z transformed correlations ± SEM) comparing activations within core language hubs (purple), peripheral regions (green), and connectivity (yellow) between these regions for GPT-2 embeddings from layers 8 and 12 during Movie (left) and Audio (right) tasks. Connectivity consistently outperformed activations alone in both tasks and across layers, emphasizing that linguistic representations emerge from extensive dynamic interactions across distributed brain networks rather than isolated activation in traditional language regions. Significant differences (*$P < 0.001$) are indicated by brackets.

**Fig. S5. Distribution of activity for words and their sensorimotor experiential associations during audiobook listening.**

The white outline corresponds to the main effect of words and the colours represent the 11 sensorimotor modulators of those same words. Specifically, the 11 sensorimotor modulators (e.g., foot/leg vs. baseline) are in red whereas the pairwise contrasts between modulators (e.g., foot/leg vs. hand/arm) are all shown in yellow. Insignificant voxels are in blue. All results are cluster-size corrected for multiple comparisons at $\alpha = 0.01$ and displayed with a minimal cluster of 20 voxels (160 μL).

**Fig. S6. Distribution of high versus low centrality regions aggregated over time during audiobook listening.**

Dynamic functional connectivity was conducted and colours show high minus low centrality regions averaged over time. Significant high > low regions that also exceed a 90% value to be considered hubs are in red. Spatial independent component and dual regression analyses were also conducted on the high and low centrality time series across participants to find centrality hubs. One component had a high correlation with the main effect of words (black outline) and was high > low centrality. The white outline corresponds to the main effect of words for comparison. All results were corrected for multiple comparisons.

*Fig. S1*

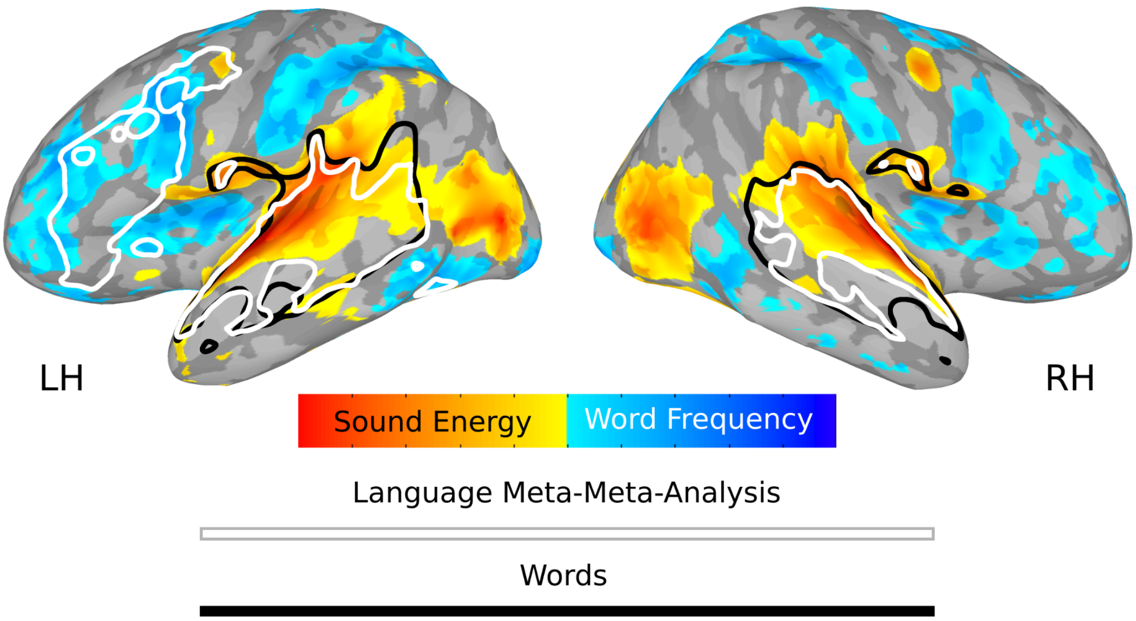
LH
RH
Sound Energy
Word Frequency
Language Meta-Meta-Analysis
Words

***Fig. S2***

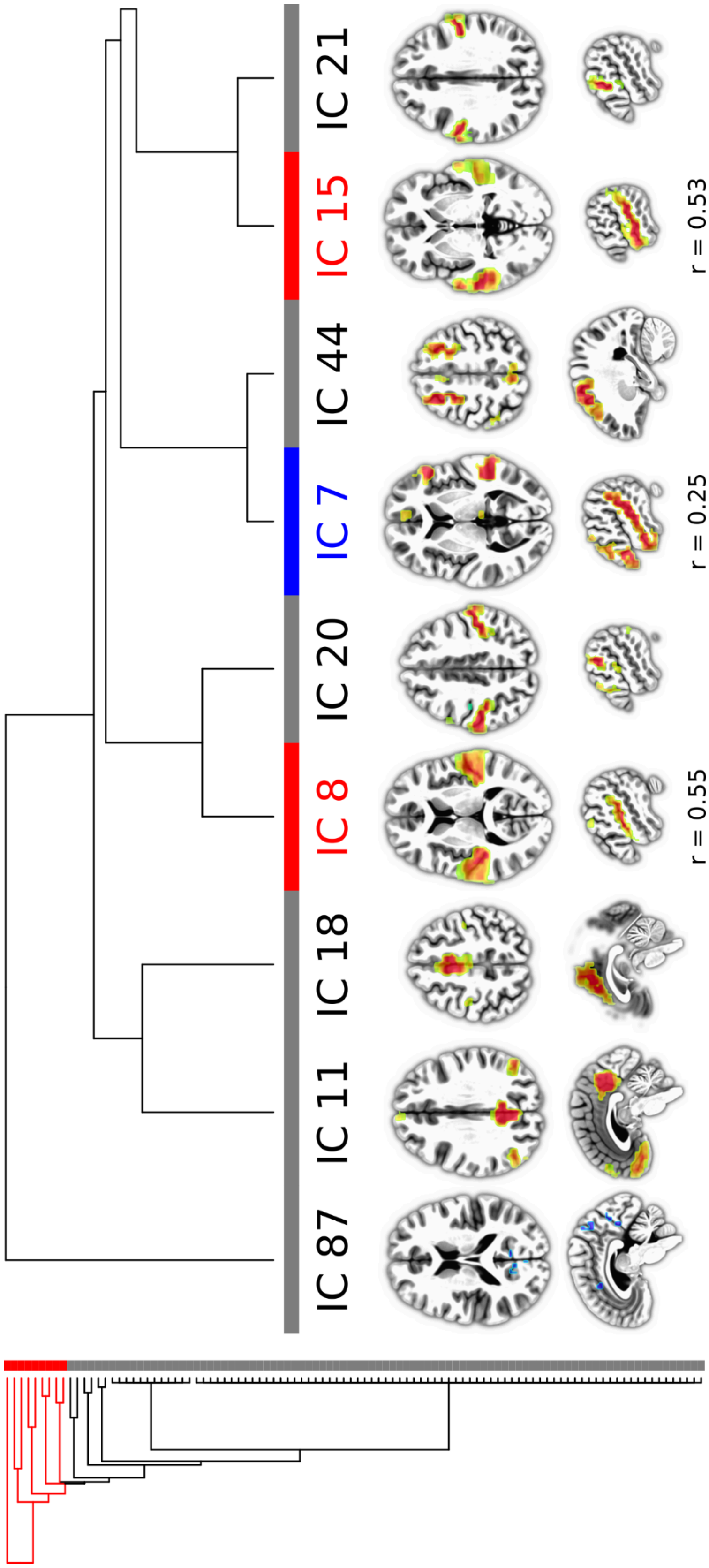

***Fig. S3***

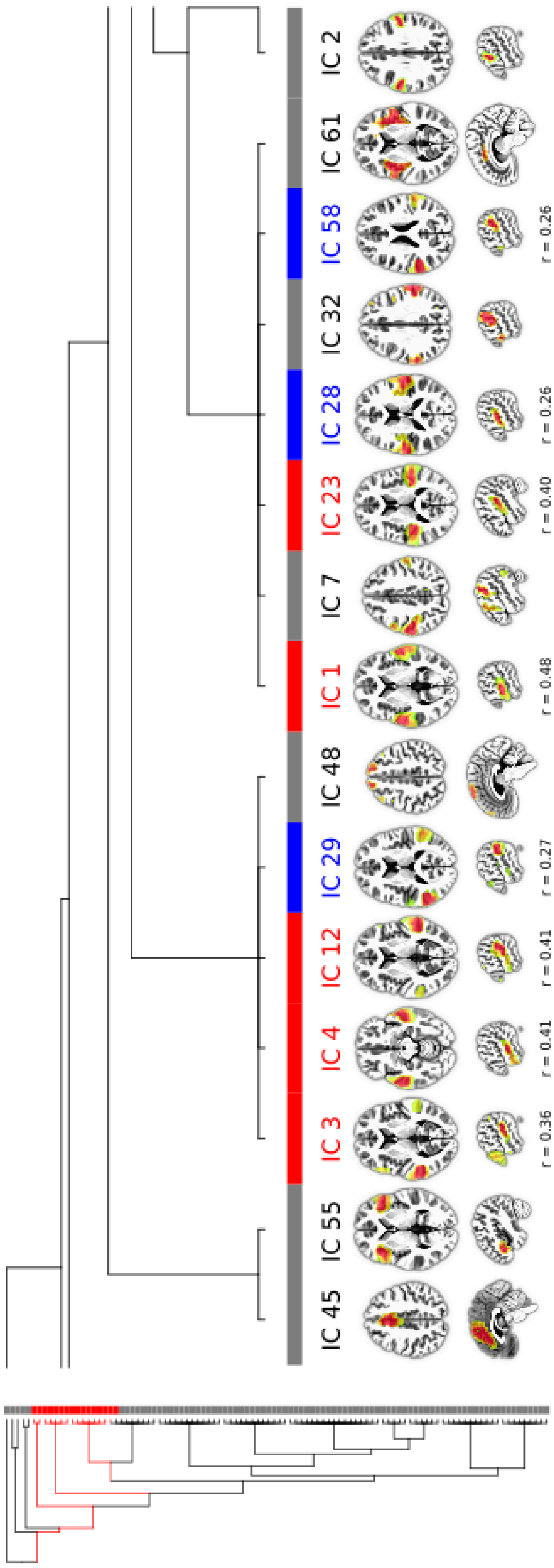
IC 2
IC 61
IC 58
r = 0.26
IC 32
IC 28
r = 0.26
IC 23
r = 0.40
IC 7
IC 1
r = 0.48
IC 48
IC 29
r = 0.27
IC 12
r = 0.41
IC 4
r = 0.41
IC 3
r = 0.36
IC 55
IC 45

***Fig. S4***

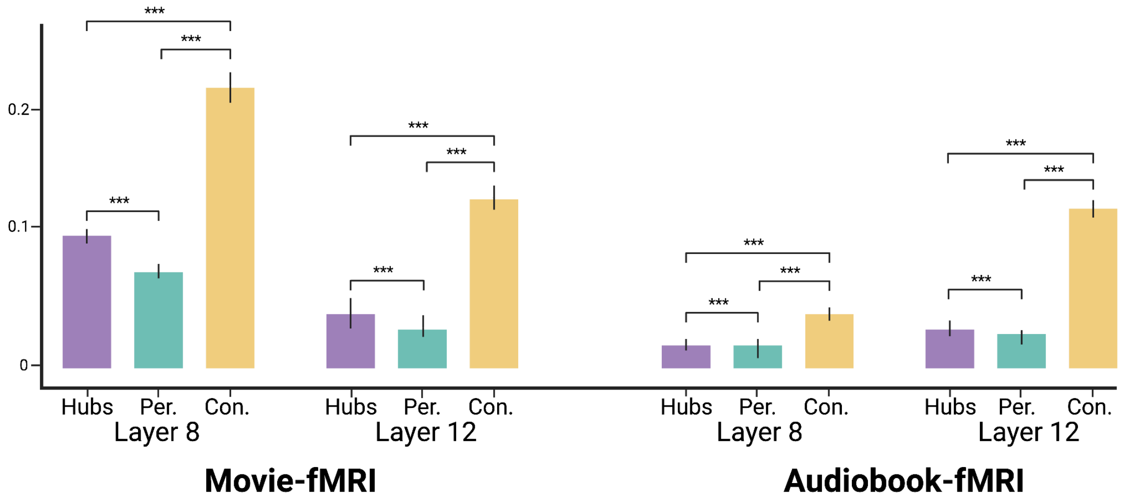
***
0.2
0.1
0
Hubs
Per.
Con.
Layer 8
Layer 12
Movie-fMRI
Audiobook-fMRI

*Fig. S5*

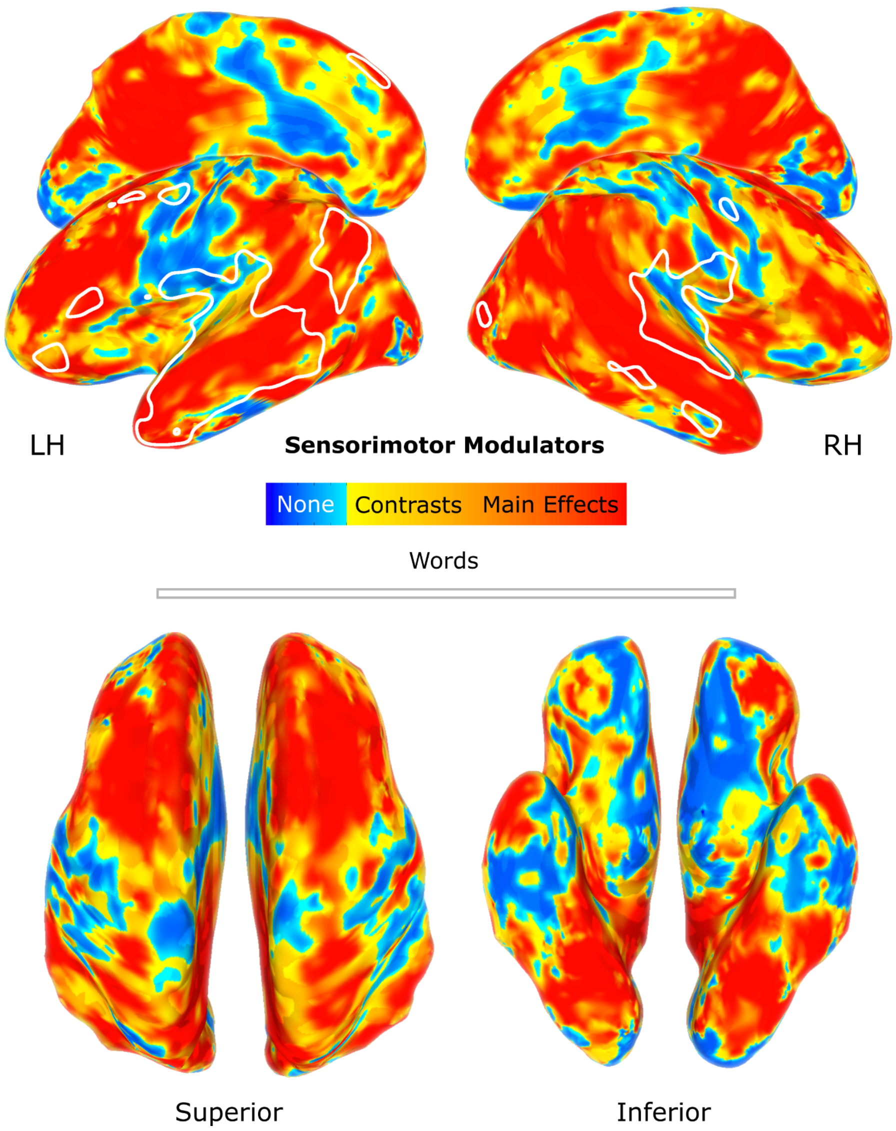
LH
Sensorimotor Modulators
RH
None
Contrasts
Main Effects
Words
Superior
Inferior

***Fig. S6***

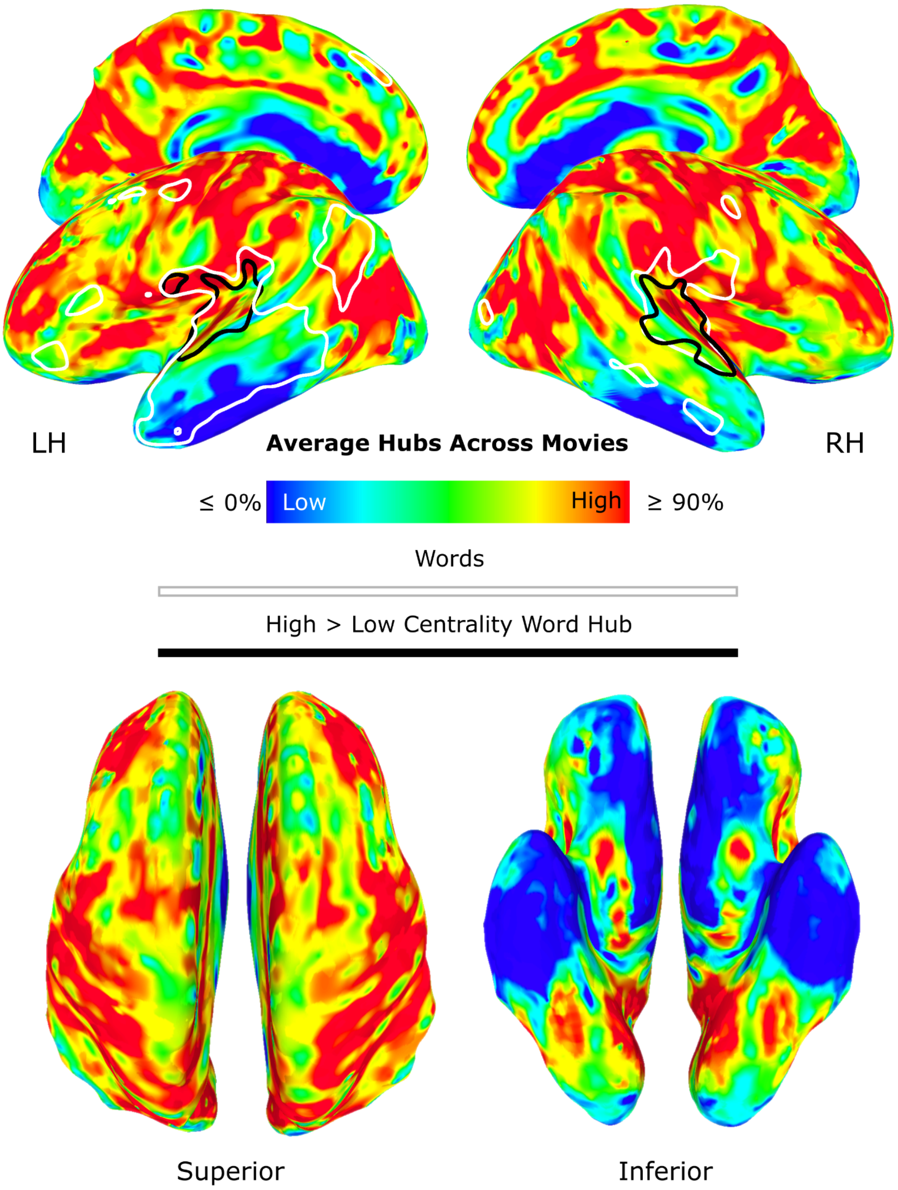
LH
Average Hubs Across Movies
RH
≤ 0%
Low
High
≥ 90%
Words
High > Low Centrality Word Hub
Superior
Inferior
